# Sample size calculation for estimating key epidemiological parameters using serological data and mathematical modelling

**DOI:** 10.1101/287581

**Authors:** Stéphanie Blaizot, Sereina A. Herzog, Steven Abrams, Heidi Theeten, Amber Litzroth, Niel Hens

**Affiliations:** Centre for Health Economics Research and Modelling Infectious Diseases (CHERMID), Vaccine and Infectious Disease Institute (VAXINFECTIO), University of Antwerp, Antwerp, Belgium; Institute for Medical Informatics, Statistics and Documentation, Medical University of Graz, Graz, Austria; Interuniversity Institute for Biostatistics and statistical Bioinformatics, UHASSELT (Hasselt University), Hasselt, Belgium; Centre for the Evaluation of Vaccination, Vaccine and Infectious Disease Institute (VAXINFECTIO), University of Antwerp, Antwerp, Belgium; Service of Epidemiology of infectious diseases, Operational Directorate Public health and surveillance, Scientific Institute of Public Health (WIV-ISP), Brussels, Belgium

**Keywords:** infectious diseases, mathematical models, study design, sample size, allocation, precision

## Abstract

**Background:** Our work was motivated by the need to, given serum availability and/or financial resources, decide on which samples to test for different pathogens in a serum bank. Simulation-based sample size calculations were performed to determine the age-based sampling structures and optimal allocation of a given number of samples for testing across various age groups best suited to estimate key epidemiological parameters (e.g., seroprevalence or force of infection) with acceptable precision levels in a cross-sectional seroprevalence survey.

**Methods:** Statistical and mathematical models and three age-based sampling structures (survey-based structure, population-based structure, uniform structure) were used. Our calculations are based on Belgian serological survey data collected in 2002 where testing was done, amongst others, for the presence of IgG antibodies against measles, mumps, and rubella, for which a national mass immunisation programme was introduced in 1985 in Belgium, and against varicella-zoster virus and parvovirus B19 for which the endemic equilibrium assumption is tenable in Belgium.

**Results:** The optimal age-based sampling structure to use in the sampling of a serological survey as well as the optimal allocation distribution varied depending on the epidemiological parameter of interest for a given infection and between infections.

**Conclusions:** When estimating key epidemiological parameters with acceptable levels of precision within the context of a single cross-sectional serological survey, attention should be given to the age-based sampling structure. Simulation-based sample size calculations in combination with mathematical modelling can be utilised for choosing the optimal allocation of a given number of samples over various age groups.

## Introduction

Several key epidemiological parameters such as the prevalence, the force of infection (i.e., the instantaneous rate at which susceptible individuals become infected) or the basic reproduction number R_0_ (i.e., the expected number of secondary cases produced by a typical infected person during his/her entire period of infectiousness when introduced into a totally susceptible population) can be computed through the use of mathematical models.

Mathematical models for infectious diseases often rely on data from serological surveys and the usefulness of these surveys has recently been highlighted.^1^ Specifically, in a cross-sectional survey, samples taken from individuals at a certain time point provide (at least partial) information about whether or not these individuals have been immunised (infected or vaccinated) before that time point (depicting current status data). In practice, antibodies which were formed in response to an infecting organism or following vaccination are identified in the serum. Typically, the antibody levels from serological data are compared to a predetermined cut-off level to determine the individuals’ humoral immunological status. Under the assumptions of lifelong humoral immunity and an epidemic in a steady state (i.e., at equilibrium), the age-specific force of infection can be estimated from such data.^2^

Publications that used mathematical modelling to inform the design of studies, including cross-sectional studies, are scarce.^3^ Moreover, only a few studies used mathematical or statistical models to inform the design of serological surveys. Marschner^4^ introduced a method for determining the sample size of a cross-sectional seroprevalence survey to estimate the age-specific incidence of an irreversible disease, based on the illness-death model assuming time homogeneity and non-differential mortality as described in Keiding’s 1991 paper.^5^ More recently, Nishiura *et al.*^6^ proposed a framework to compute the uncertainty bounds of the final epidemic size to H1N1-2009 and to determine the minimum sample size required. Sepúlveda and Drakeley^7^ proposed two sample size calculators, depending on whether the seroreversion rate (i.e., rate of antibody decay) is known, for estimating the seroconversion rate in malaria transmission in low endemicity settings using a reverse catalytic model. They extended the method to determine the sample size required to detect a reduction in the seroconversion rate at a given time point before survey sampling caused by a field intervention.^8^ Lastly, Vinh and Boni^9^ assessed the power of serial serological studies in inferring key epidemiological parameters using a mathematical model.

In this paper, simulation-based sample size calculations are performed in order to determine the age-based sampling structures and optimal allocation distributions best suited to estimate key epidemiological parameters with acceptable precision levels. Specifically, we use four models and three age-based sampling structures within the context of a single cross-sectional seroprevalence survey. We differentiate between endemic and non-endemic settings. In the latter case, we limit ourselves to estimating the prevalence and defer extensions thereof to future work. The objectives of this paper are i) to investigate to what extent the precision of key epidemiological parameters is modified in these models when assuming different age-based sampling structures, thereby giving insights into the optimal age structure; ii) to provide an order of magnitude of the sample size required to attain a specified precision for a particular parameter; and iii) to give insights into the optimal allocation of a fixed sample size among age groups.

Our work is motivated by the need to, given serum availability and/or financial resources, decide on which samples to test for different pathogens in a serum bank.

## Methods

### Data

A serological survey testing for the presence of, amongst others, measles, mumps, rubella, varicella-zoster virus (VZV), and parvovirus B19 IgG antibodies was conducted on large representative national serum banks in Belgium. The sera were collected between 2001 and 2003 and were obtained from residual sera submitted for routine laboratory testing (individuals aged <18 years) or from blood donors (18 years and over). This survey was designed as proposed by the European Sero-Epidemiology Network (ESEN) which aimed to standardize the serological surveillance of immunity to various diseases in European countries.^10^ In particular, children and adolescents were oversampled in the survey. A total of 3378 samples were collected and the age of the individuals ranged from 0 to 65 years. The number of samples with immunological status with regard to measles, mumps, rubella, VZV, and parvovirus B19 infection were 3190, 3004, 3195, 3256, and 3080, respectively. Samples from children aged less than 6 months or 1 year were omitted in our analyses because of distortions expected from the presence of maternal antibodies against the various pathogens. Since a national immunisation programme against measles, mumps, and rubella has been introduced in 1985 in Belgium with gradually increasing vaccine coverage in the targeted age groups (infants, adolescents aged 11-13 years, and catch-up campaigns in adults), endemic equilibrium for these infections in 2002 cannot be assumed. In constrast, no immunisation programme against VZV and parvovirus B19 has been introduced, making endemic equilibrium a tenable assumption for both infections.

Ethical approval for the setup of the 2002 serum set was obtained from the Ethics Committee of the University of Antwerp.

### Models

Here, we briefly present an overview of the methods used to derive key epidemiological parameters from serological survey data and we refer to Hens *et al.*^2^ for a more in-depth explanation of the methodology. We start from the basic concept of an age-specific prevalence and gradually move to the force of infection and other key epidemiological parameters such as the basic and effective reproduction numbers in endemic equilibrium.

Age-specific seroprevalence can be modelled in the framework of generalized linear models (GLMs; see e.g. Hens *et al*.^11^). For estimating the (age-specific) force of infection from seroprevalence data, various statistical methods have been used in the literature including linear and non-linear parametric (e.g., fractional polynomials or catalytic model) and non-parametric approaches. Complementarily, the flow of individuals between the mutually exclusive stages of an infectious disease can be described using compartmental dynamic transmission models. The simplest such model, the SIR model, describes the flow between the susceptible (*S*), the infected and infectious (*I*), and recovered class (*R*). The following set of partial differential equations in continuous age and time can be used to describe the SIR dynamics mathematically:

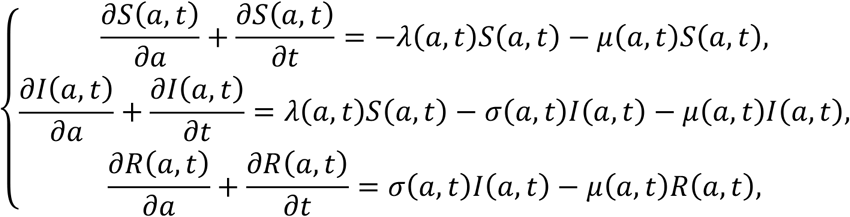

with the age- and time-specific population size given by *N* (*a,t*) = *S* (*a,t*) + *I* (*a,t*) + *R* (*a,t*) and with *λ* (*a,t*) the force of infection, *σ* (*a,t*) the recovery rate, and *μ* (*a,t*) the all-cause mortality rate.

Assuming a closed population of size *N* in demographic and endemic equilibrium, we obtain a set of ordinary differential equations (ODEs):

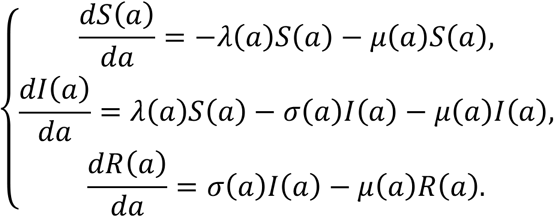

Solving the above set of ODEs, the following expression for the seroprevalence of individuals of age, *a* is obtained:

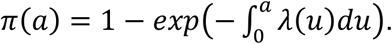

It is possible to solve the above equation numerically by turning to a discrete age framework, thereby assuming a constant force of infection *λ*_*k*_ in each age class [*a*_[*k*]_, *a*_[*k*.1]_], *k* = 1, …, *J*.^12^ The seroprevalence at age *a* in the *j*^th^ age interval is approximated by:

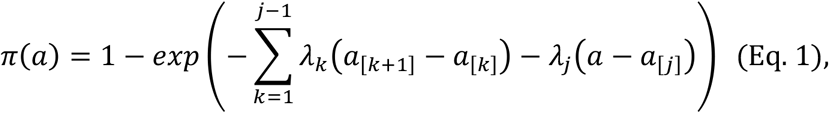

where *a*_[1]_ = 0 and *a*_[*J*.1]_ = *L* (the life expectancy).

From this model, other key epidemiological parameters can be calculated such as the basic and effective reproduction number (R_0_ and R_eff_ respectively; R_eff_ reflects the actual average number of secondary cases that can be observed in a partially immune population) or the average age at infection.

Since seropositive results for measles, mumps, and rubella are a mix of vaccine- and infection-induced immunity, implying time-heterogeneity which is beyond the scope of this paper, only the age-specific seroprevalence for these diseases was modelled. We considered a logistic model with piecewise constant prevalence values within the following age classes based (partially) on vaccination policies: [1,2), [2,11), [11,16), [16,21), [21,31), and [31,65] years. The estimates of the coefficients using this model (on the logit scale) are denoted by 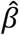.

For infections under endemic equilibrium, three mathematical models for estimating the force of infection were considered. The first is an MSIR model with piecewise constant force of infection which is a slight adaptation of the model in (Eq. 1). The seroprevalence at age *a* in the *j*^th^ age interval is approximated by:

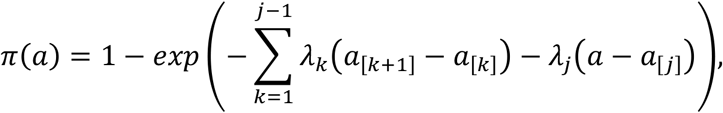

with *a*_[1]_ = *A*, where *A* is the age at which maternal immunity is lost. In this paper, we considered an MSIR model with piecewise constant force of infection within the following six age classes based on school enrollment ages in Belgium (except for the oldest age group): [1,2), [2,6), [6,12), [12,19), [19,31), and [31,65] years.

The second model considered in this paper is the exponentially damped model for the force of infection as described by Farrington.^13^ This model assumes that the force of infection increases to a peak in a linear fashion followed by an exponential decrease, and can be formulated as follows:

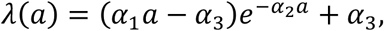

with *α*_1_, *α*_2_ and *α*_3_ the model parameters to be estimated from the data. Integrating *λ*(*a*) results in a non-linear model for the seroprevalence, i.e.,

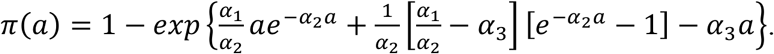

We considered a third model for parvovirus B19 infection, a mathematical model allowing for boosting and waning immunity, since lifelong protection against infection upon recovery from parvovirus B19 is questionable.^14–17^ Goeyvaerts *et al*.^18^ considered several extensions of the MSIR model to account for waning of disease-acquired antibodies and/or for boosting of low immunity by exposure to infectious individuals. Here, we used the model with the best Akaike information criterion (AIC) value which was the compartmental model allowing for age-specific waning of disease-acquired antibodies and boosting of low immunity, denoted by “MSIRWb-ext AW” (see the Supplementary Material). In this model, individuals move from a high immunity state R to a low immunity state W at a rate *ϵ*_1_ and *ϵ*_2_ for age group <35 and ≥35 years respectively. In addition, low immunity can be boosted by exposure to infectious individuals; the boosting rate was assumed to be proportional to the force of infection by a factor of *ϕ*. The transmission rates are assumed to be directly proportional to age-specific rates of making social contact with a proportionality factor *q*. To be consistent with the aforementioned paper, only the samples from children aged less than 6 months were omitted for this analysis.

The first two columns of Table 1 show a summary of the models used for each of the pathogens studied. Formulas to calculate the key epidemiological parameters (i.e., age-standardized seroprevalence and force of infection, R_0_, R_eff_, and the average age of infection) can be found in the Supplementary Material. The age-specific sero-prevalence and force of infection were calculated in the following age groups: [1,2), [2,6), [6,12), [12,19), [19,31), and [31,65] years.

**Table 1.**
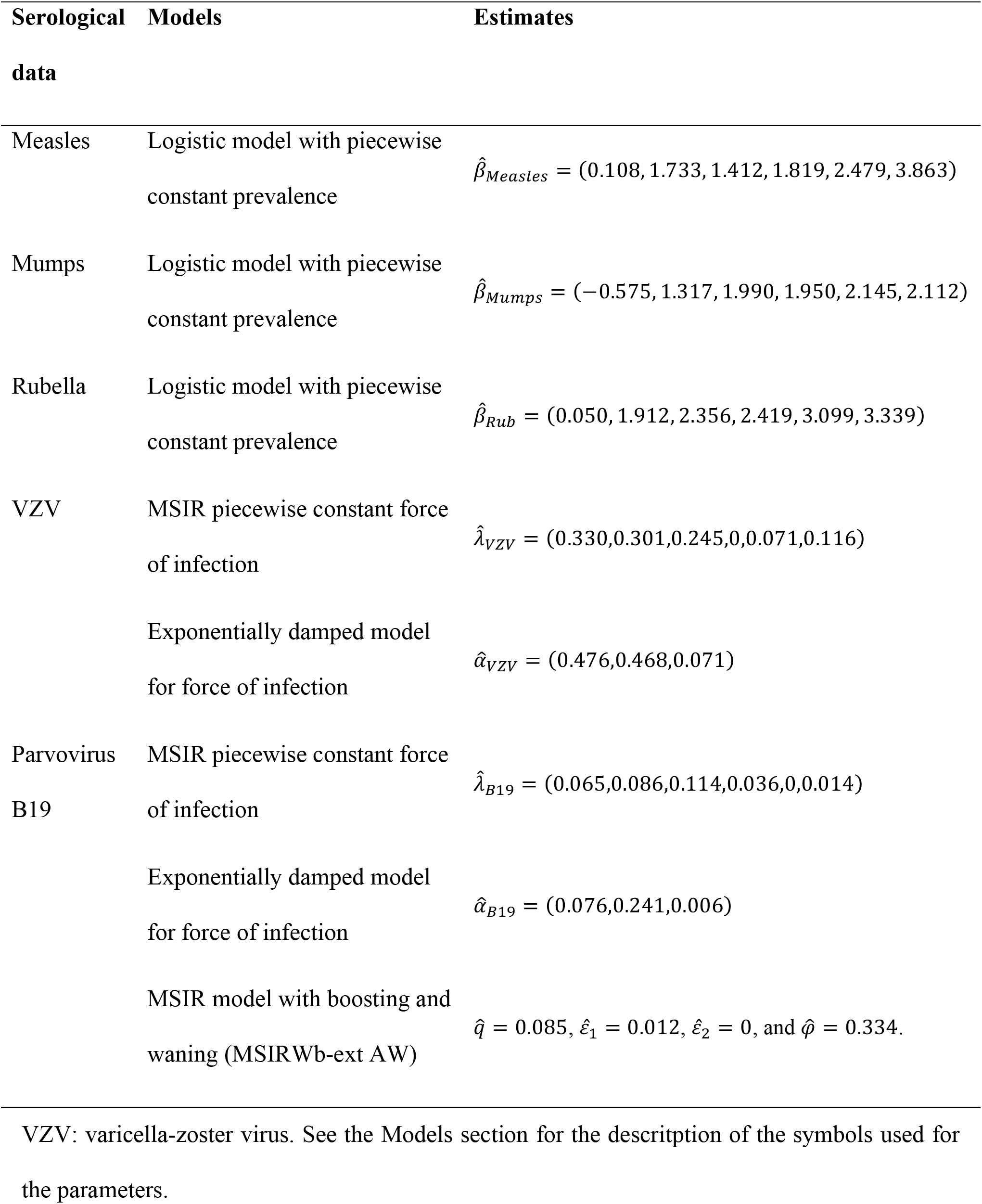
Summary of the models considered for each of the pathogens and the corresponding model parameter estimates using the observed serological survey data

### Estimating the model parameters

Maximum likelihood estimates were obtained for each model and pathogen assuming that the observed prevalence follows a binomial distribution. The analyses were performed using R software (version 3.3.1).^19^ Using the estimated values of the parameters for each model and pathogen (with age values rounded down to integer values), age-specific “true” prevalence values were calculated which were used in the simulations (see next section).

### Simulations

Figure 1 gives a schematic representation of the approach used in this paper. Three age structures were compared: the age structure derived from the pathogen-specific data of the serological survey in which children and adolescents were oversampled (survey-based), the age structure of the Belgian population in 2003 (population-based),^20^ and a uniform age structure (see Supplementary Material: Figure S1 and Table S1). In order to calculate the (simulation-based) precision of the estimates of the epidemiological parameters, 500 datasets were generated for each model using a binomial distribution with model- and age-specific “true” prevalence values and age-specific sample sizes. These age-specific sample sizes depend on the age structure and the total sample size (N=1650, 3300, 6600, 9900, 13200, or 19800) that are used. The precision around the estimated key epidemiological parameters is defined to be half the length of the 95% percentile-based confidence interval (CI) calculated over the 500 simulations. In the MSIR model with piecewise constant force of infection for the VZV infection, simulations with biologically implausible estimated values (>10) were excluded; such values were obtained in the age group >30 years due to a simulated prevalence of 100% in this age group. These simulations were replaced. Here, the simulations will give insights into the age structure best suited to estimate given key parameters but also provide insights into the sample size needed.

**Figure 1.**
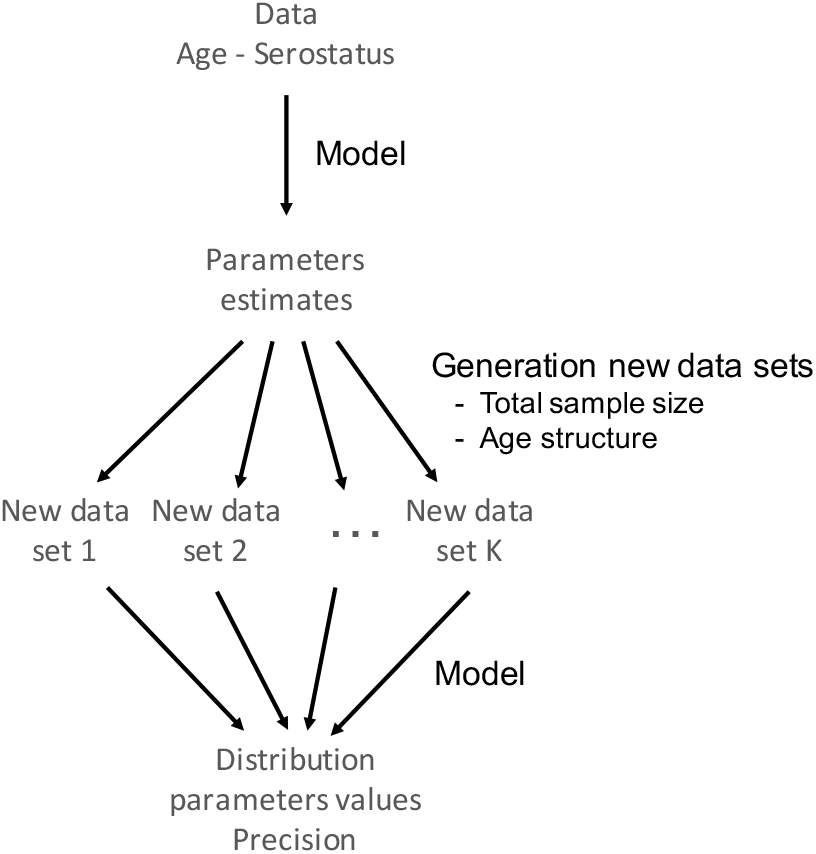
Schematic representation of the approach used in this paper.

Alternatively, if resources are available for a predetermined number of samples or if one wants to test samples previously collected without prior knowledge of age-based sampling structure, the proportion of the samples to allocate in each age group could be investigated to obtain the highest precision for a given parameter. Here, the optimal allocation was determined by calculating the precisions obtained using different distributions. To restrict the number of distributions to compare, we varied the proportions among the six age groups ([1,2), [2,6), [6,12), [12,19), [19,31), and [31,65] years) from 10% to 50% (leading to 126 distributions) and assuming a uniform distribution within each age group. We investigated the optimal allocation for several values of the total number of samples available: N= 1650, 3300, 6600, 9900, 13200, or 19800. Five hundred datasets were generated for each distribution and each sample size. For the seroprevalence and force of infection by age group, the age distribution providing the best joint precision, defined as the sum of the precisions in each age group, is reported.

## Results

### Estimates of the model parameters obtained using the observed serological survey data

The model estimates for each of the different pathogens are given in Table 1. Figure S3 (Supplementary Material) shows the estimated prevalence and force of infection for each model and disease. The results between the models were close; however, as expected, for parvovirus B19, the MSIRWb-ext AW model was able to capture, though only slightly, the decrease in seroprevalence around age 30. In this model, the force of infection had a bimodal shape (with modes around ages 7 and 35 years; Figure S3).

Since our simulations were based on integer age values, the MSIR and MSIRWb-ext AW models were re-run after rounding continuous age values down to integers; however, the estimates were close when using continuous or integer values. The estimates obtained using the MSIR model with piecewise constant force of infection were: 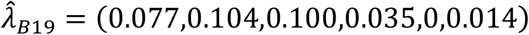, 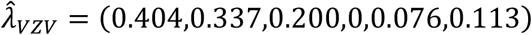. The following estimates were obtained using the MSIRWb-ext AW model for parvovirus B19: 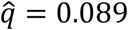, 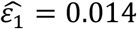, 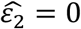, and 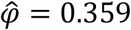. Estimates of the key epidemiological parameters are provided in the Supplementary Material (Tables S2-S4).

### Comparisons of the three age-based sampling structures

For the overall seroprevalence of measles and VZV, in both models used, the survey-based age structure led to the best precision (Figures 2 and 3, Table S5, Tables S8-S9). However, when modelling mumps and parvovirus B19, in the three models used, the precision of the overall seroprevalence was found to be better using a uniform or population-based age structure (Figures 2 and 4, Table S6, Tables S10-12). Finally, the precision for the estimated overall rubella seroprevalence was similar for the three different age structures (Figure 2, Table S7).

**Figure 2.**
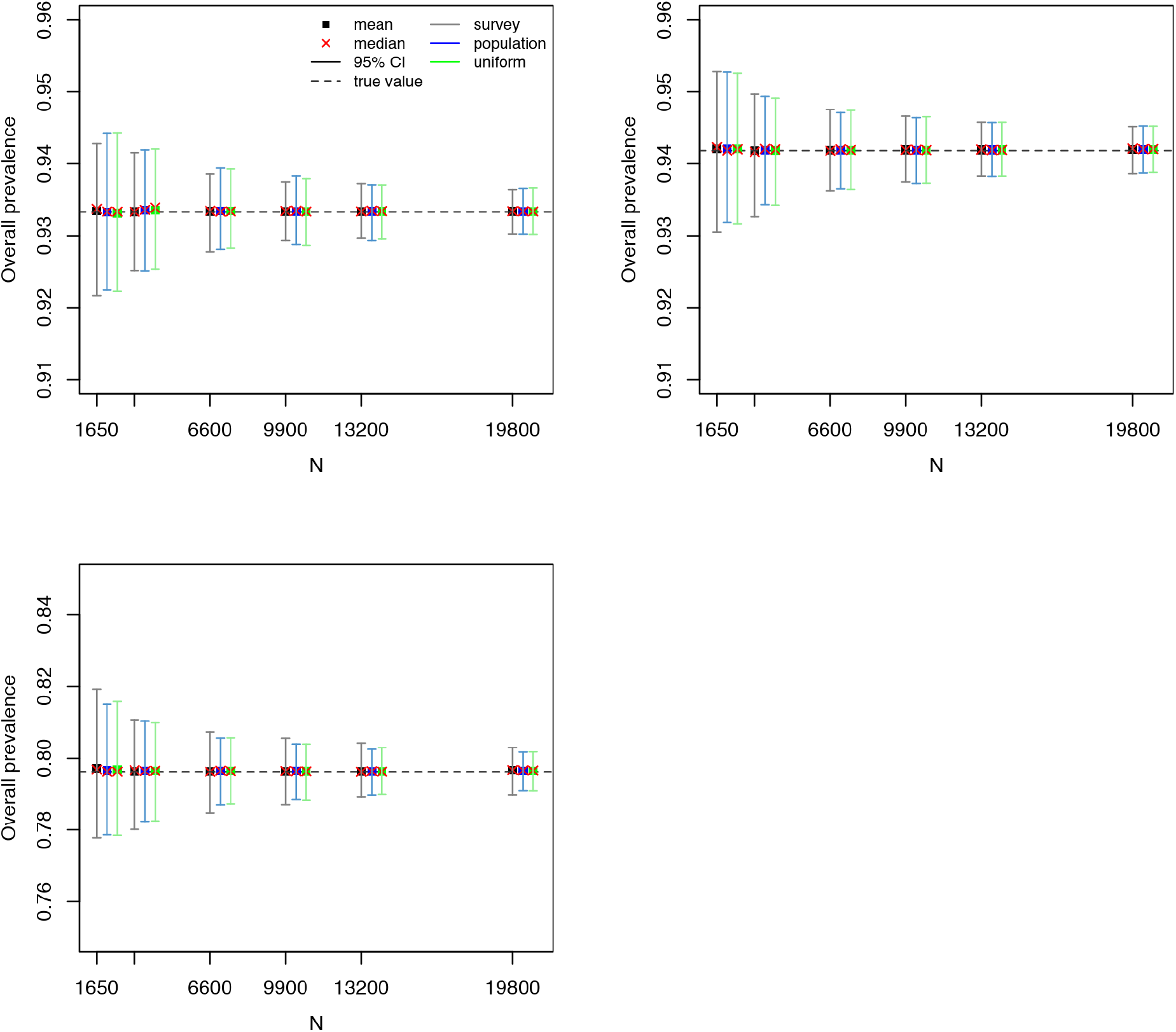
Measles, mumps, rubella serological data: mean, median, and 95% confidence interval for the overall seroprevalence over 500 simulations as a function of the total number of sampled individuals (N) using the logistic model with piecewise constant prevalence. Top left: Measles. Top right: Rubella. Bottom: Mumps. “True” overall seroprevalence is the estimated overall seroprevalence using the models on the observed serological survey data (with integer age values). The y-axes have different ranges of values for better legibility.

**Figure 3.**
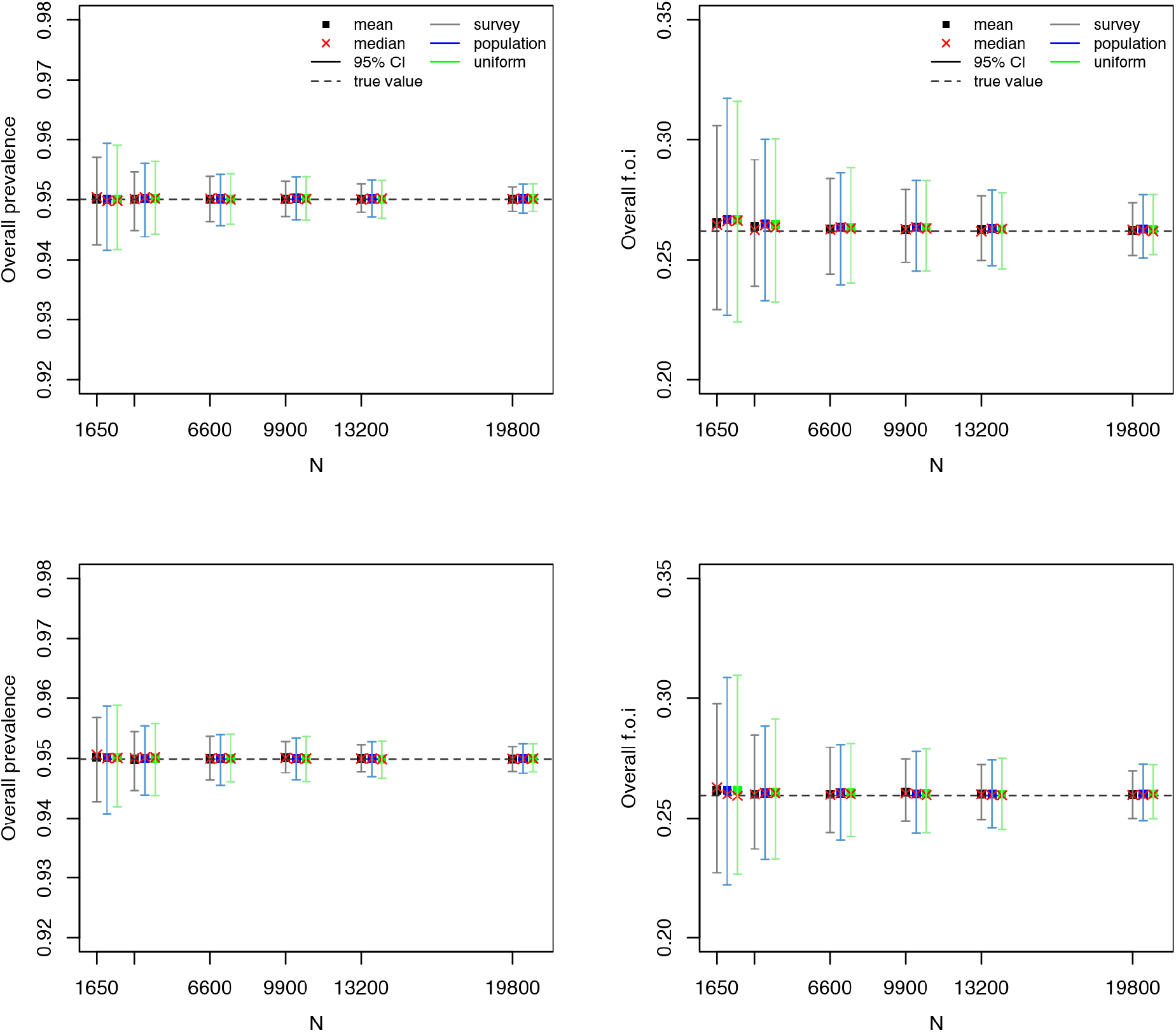
VZV serological data: mean, median, and 95% confidence interval for the overall seroprevalence (left) and overall force of infection (right) over 500 simulations as a function of the total number of sampled individuals (*N*) for the MSIR model with piecewise constant force of infection (top) and the exponentially damped model (bottom). “True” overall seroprevalence is the estimated overall seroprevalence using the models on the observed serological survey data (with integer age values).

**Figure 4.**
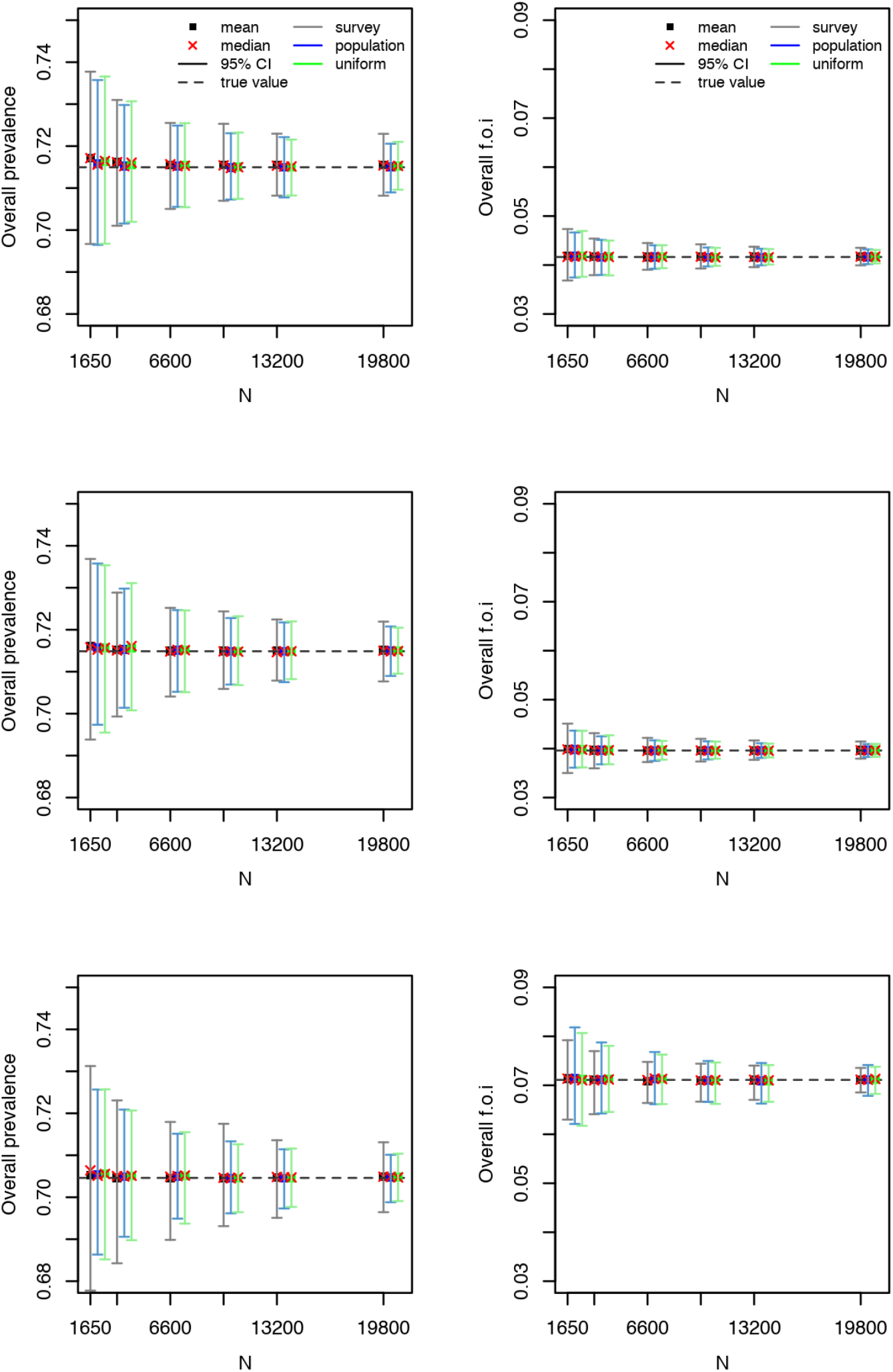
Parvovirus B19 serological data: mean, median, and 95% confidence interval for the overall seroprevalence (left) and overall force of infection (right) over 500 simulations as a function of the total number of sampled individuals (*N*) for the MSIR model with piecewise constant force of infection (top), the exponentially damped model (middle), and the MSIRWbext-AW model (bottom). “True” overall seroprevalence is the estimated overall seroprevalence using the models on the observed serological data (with integer age values).

The precision of the estimated overall force of infection was better when using the survey-based age structure for VZV infection, in both models used (Figure 3, Tables S8-9) and for parvovirus B19 infection under the MSIRWb-ext AW model, and using a uniform or population-based age structure for parvovirus B19 infection in the two other models used (Figure 4, Tables S10-12). For all the pathogens, as could be expected given the oversampling in children and adolescents in the survey-based age structure, the precision of the estimated seroprevalence by age group was better when using the survey-based age structure in the young age groups and the uniform or population-based age structure for the oldest age groups (see Tables S5-S12 in the Supplementary Material). The same pattern was observed for the force of infection of VZV and parvovirus B19 by age group (see Tables S8-S12 in the Supplementary Material).

In the exponentially damped model, the precision of R_0_ and the average age at infection was slightly better using the uniform or population-based age structure for parvovirus B19 while it was better using the survey-based age structure for VZV (see Tables S8 and S10). In the MSIRWbext-AW model, the precision of R_0_, R_eff_, and the average age at infection of parvovirus B19 was slightly better using the survey-based age structure while that of the relative boosting factor (*ϕ*) was better using the uniform or population-based age structure (Figure 5 and Table S12). However, the precision of this factor was poor, with large confidence intervals, and the average age at infection should be interpreted with caution given the bimodal force of infection.

**Figure 5.**
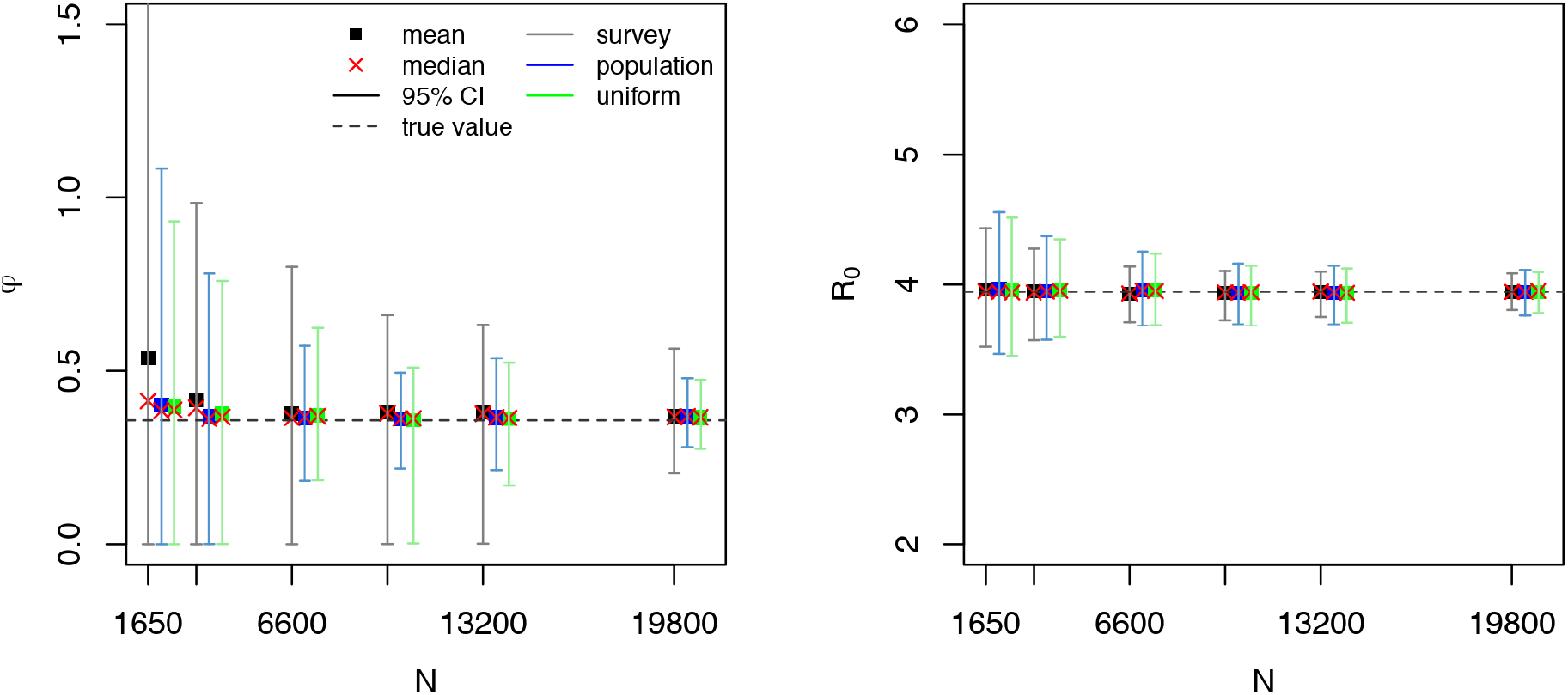
Parvovirus B19 serological data: mean, median, and 95% confidence interval for the relative boosting factor *ϕ* (left) and basic reproduction number *R*_0_ (right) over 500 simulations as a function of the total number of sampled individuals (*N*) for the MSIRWbext-AW model. “True” value is the value estimated using the model on the observed serological data (with integer age values). The y-axes have different ranges of values for better legibility.

### Sample size needed

To obtain a 2% precision around the overall seroprevalence estimate, the sample size needed would be around 1650 for mumps and parvovirus B19, while a lower number of samples would be sufficient for measles, VZV, and rubella; to obtain a 1% precision the sample size needed would be around 6600 for mumps and parvovirus B19, and 1650 for measles, VZV, and rubella (Figures 2-4; Tables S5-S12). These results were quite consistent across age structures.

### Optimal allocation of a fixed sample size among age groups

For the overall seroprevalence of measles, mumps, or rubella, the optimal allocation (distribution over age groups) of a fixed number of samples would be a distribution with a high percentage of the data among age groups [19,31) and [31,65] years, for each sample size used (Table S13-S15 in the Supplementary Material). Regarding the seroprevalence by age group, for measles, mumps and rubella, we have noticed some variations across the sample sizes; the optimal allocations were broadly uniform across the age groups.

The optimal allocation for overall VZV seroprevalence or force of infection estimates varied with sample size; the oldest two age groups would rather be favoured (Figure 6 and Table S16-S17 in the Supplementary material). The optimal allocation for the overall parvovirus B19 seroprevalence estimate would be a distribution with a high percentage of data in the oldest age group, for each model and sample size used (Figure 7 and Table S18-S20 in the Supplementary Material). Regarding the overall force of infection of parvovirus B19, the optimal allocation would entail a distribution with high percentage among the oldest age group in the MSIR model with piecewise constant force of infection and exponentially damped model, while more equally distributed over the various age groups for the MSIRWb-ext AW model.

**Figure 6.**
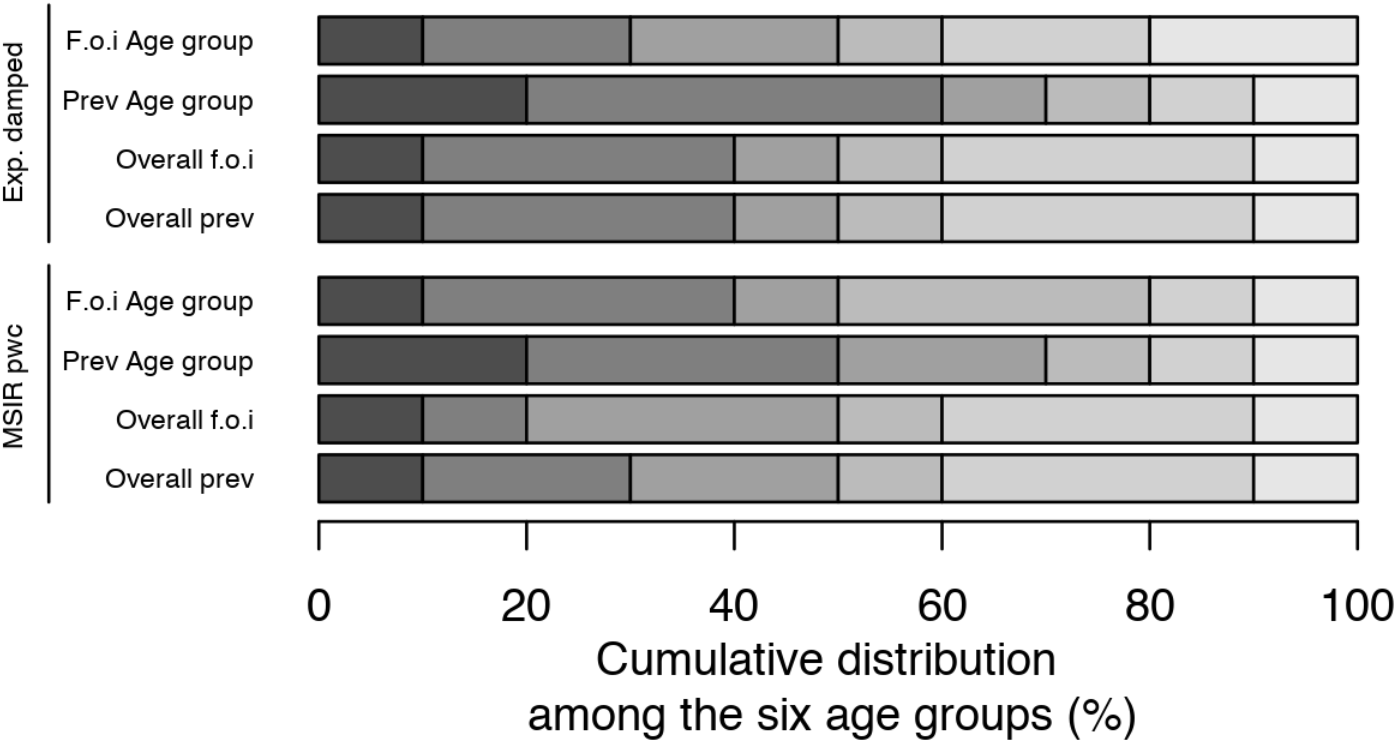
VZV serological data: optimal allocation (N=3300) for various key epidemiological parameters and by model (y-axis) among the six age groups (with lighter shades with increasing age group): [1,2), [2,6), [6,12), [12,19), [19,31), and [31,65].

**Figure 7.**
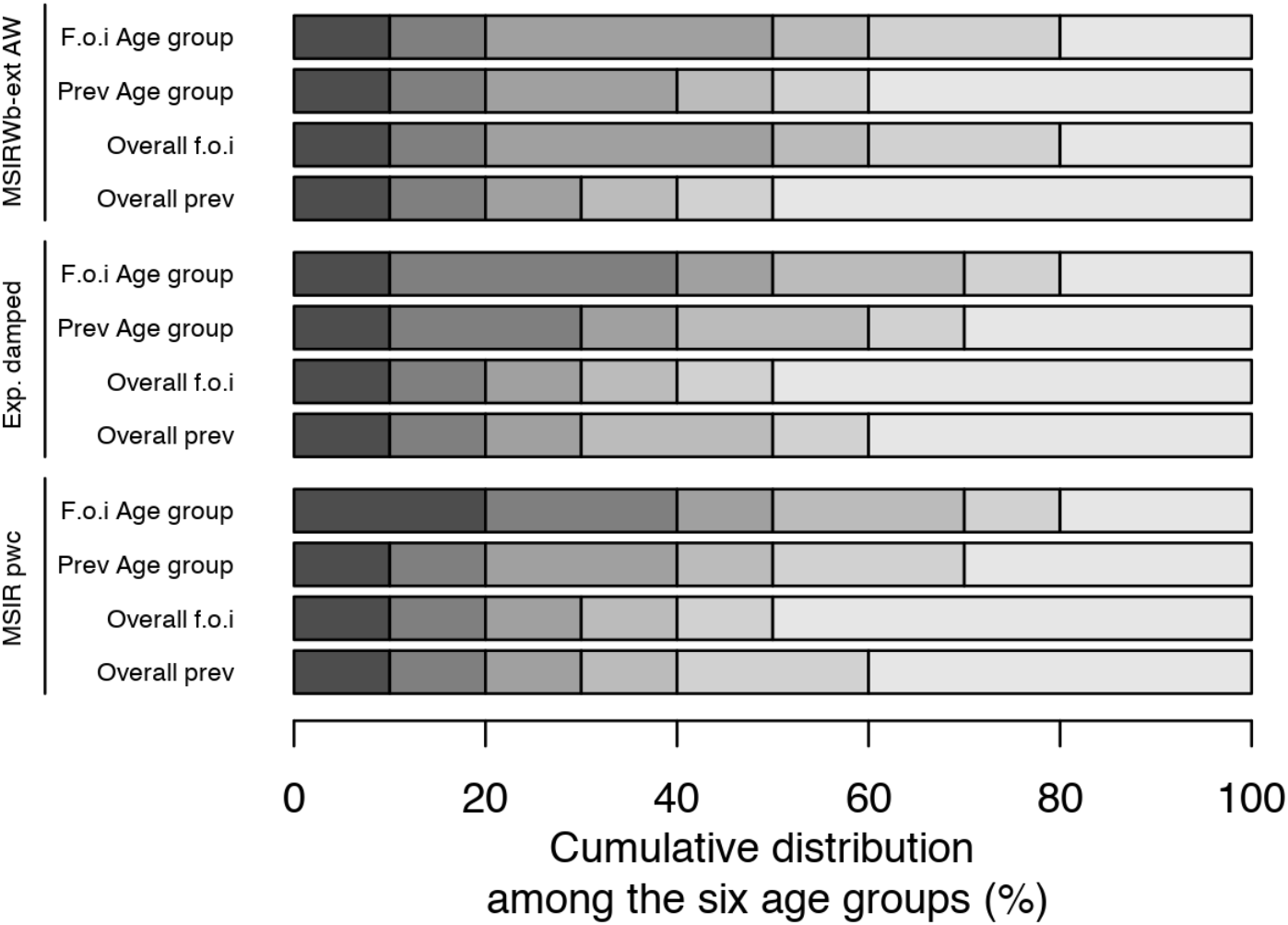
Parvovirus B19 serological data: optimal allocation (N=3300) for various key epidemiological parameters and by model (y-axis) among the six age groups (with lighter shades with increasing age group): [1,2), [2,6), [6,12), [12,19), [19,31), and [31,65].

Regarding the seroprevalence or force of infection by age group for VZV and parvovirus B19, some variations between models and sizes were observed; the optimal allocations were broadly uniform across the age groups.

## Discussion

Considering sample size and optimal allocation is essential since efficient usage of resources is needed in the context of limited human or financial resources and/or time constraints for performing a serological survey. Since analytical formulas for complex models are not available, simulation-based analyses are a flexible alternative to address these considerations. In this paper, we proposed a simulation-based approach for sample size and age structure considerations, and optimal allocation of resources, in order to estimate key epidemiological parameters with acceptable levels of precision within the context of a single cross-sectional serological survey.

Our results showed that the best age structure to use in the sampling of a serological study as well as the optimal allocation distribution varied with the epidemiological parameters of interest. To our knowledge, only a few studies investigated, using mathematical or statistical models, the optimal allocation of a given number of samples over age groups to obtain good precision. Marschner^4^ showed, using an example of measles infection, that a uniform age distribution should not be optimal to obtain a good joint precision of the force of infection.

For all the infections investigated, due to the oversampling of individuals under 20 years old in the serological survey purposefully, the precision of the estimated seroprevalence by age group was better with the survey-based age structure in the young age groups and the uniform or population age structure for the oldest age groups. Moreover, because of the formulas used to compute the basic or effective reproduction number and the average age at infection, the age structure best suited to estimate these parameters was related to that of the prevalence in the exponentially damped model and of the force of infection in the MSIRWb-ext AW model. In case the boosting rate is of interest, sufficiently sampling adults is essential. Anyway, the precision of this rate was poor as was also observed in previous analyses.^18^ This could be explained by the complexity of the model used.

An important finding was that the age-specific prevalence profile, and thus the age-specific force of infection profile, had an effect on the optimal age structure to use in a serological survey or the optimal allocation for estimating the overall seroprevalence. Indeed, the optimal age structure varied between VZV and parvovirus B19 infections, the seroprevalence increasing more sharply between ages 1 and 10 for VZV compared to parvovirus B19.

Our analyses could be extended to power analyses in the context of hypothesis testing. Indeed, data sets could be simulated assuming that an alternative hypothesis is true, then tested against the null hypothesis to calculate the proportion of simulated data sets in which the null hypothesis is rejected, thereby providing an estimate of the statistical power.

Other possible extensions are related to non-endemic settings. An endemic equilibrium cannot be assumed for vaccine-preventable infections such as measles, mumps, and rubella for which a national immunisation programme is in place. In such settings, dynamical mathematical models allowing time considerations could be used to calculate the sample size needed for estimating time-varying parameters with acceptable precision levels or to perform power calculations to detect changes in parameter values over time, but this needs to be investigated. In particular, these analyses could make use of serial seroprevalence surveys (i.e., repeated collections of cross-sectional population-representative serological samples).^9^ Finally, our analyses could also be extended to more complex models, for example transmission models incorporating the presence of individual heterogeneities.^21,22^

Our analyses had some limitations. First, a limited number of 500 datasets were generated to estimate the precisions in the sample size and optimal allocation considerations. However, similar results were obtained when generating 1000 or 1500 datasets (data not shown).

Second, the number of age groups to optimally allocate a given number of samples had to be limited to avoid a huge number of combinations. Here, six age groups were used leading to 126 distributions. Alternative age groups of interest or a predetermined age distribution (e.g., derived from previous surveys or population-based) can be used. Moreover, the optimal allocation will depend on the rule used to calculate the joint precision. Here, we used the sum of the age-specific precisions. Alternative rules could be considered such as the sum of the relative precisions. However, favouring very small values could result in a very large sample size or be of less interest (e.g., if force of infection in older age groups is known to be small).

Third, the use of measurements of antibody levels based on diagnostic tests relies on the assumption of a perfect test (i.e., both sensitive and specific). In lack of which, due to misclassification, the seroprevalence is not exactly equal to the disease prevalence, which would alter the estimates of the overall and age-specific prevalence, even more if sensitivity and specificity vary with age.^23^ The estimate of the seroprevalence can be corrected if estimates of the sensivity and specificity of the test(s) applied are available.^24^ Alternatively, mixture modelling of continuous antibodiy titers can be used, however the combination of this technique with mathematical models needs further investigations.^2,25-27^ In the current work, considering misclassifications negligible appeared reasonable.

Finally, like other standard methods, the approach presented here would require prior knowledge about parameter values: e.g., (sero)prevalence or force of infection by age (group) to simulate data. However, sensitivity analyses may be performed to assess how this prior knowledge affects the sample size needed or optimal allocation and would inform about the minimum sample size needed.

In any case, the choice of sampling design or modelling approach should be adapted to prior knowledge about the infection and the precision of estimates (overall or age-specific) should be considered in the context of the study goals and the anticipated implications for infection control measures or vaccine programs.

The main conclusions from the presented analyses are that attention should be given to the age-based sampling structure when estimating key epidemiological parameters with acceptable levels of precision within the context of a single cross-sectional serological survey, and that simulation-based sample size calculations in combination with mathematical modelling can be utilised for choosing the optimal allocation of a given number of samples over various age groups.

## Acknowledgements

Authors SB, HT and NH acknowledge support of the Antwerp Study Centre for Infectious Diseases (ASCID) at the University of Antwerp. NH gratefully acknowledges the chair in evidence-based vaccinology sponsored by a gift from Pfizer.

## List of Supplemental Digital Content

Supplemental Digital Content 1. SupplementaryMaterial.pdf

