## Supplementary Materials for "Sample size calculation for estimating key epidemiological parameters using serological data and mathematical modelling"

### Supplementary material

### Age structures

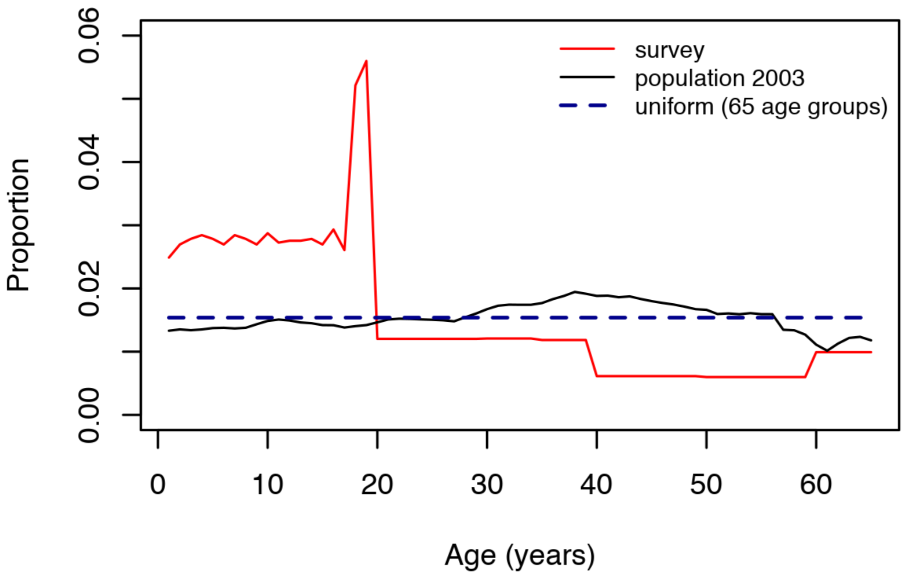

**Figure S1.** Comparison of the three age distributions.

"survey" (red line) refers to the age structure derived from the serological survey data (all individuals included, aged from 1 to 65 years old); "population 2003" (black line) refers to the age structure of the Belgian population in 2003; "uniform" (blue dashed line) refers to a uniform age structure based on 65 age groups (1-year interval).

**Table S1.** Distribution of the population among age groups by age structure

| | **Age groups** | | | | | |
|---|---|---|---|---|---|---|
| | 1 year | 2-5 years | 6-11 years | 12-18 years | 19-30 years | 31-65 years |
| **Population** | 1.1% | 4.5% | 7.2% | 8.4% | 15.3% | 63.5% |
| **Uniform** | 1.5% | 6.2% | 9.2% | 10.8% | 18.5% | 53.8% |
| **Serological survey** | | | | | | |
| Measles | 2.3% | 11.1% | 16.5% | 20.9% | 18.5% | 30.6% |
| Mumps | 2.5% | 11.3% | 16.3% | 21.6% | 18.6% | 29.7% |
| Rubella | 2.5% | 11.2% | 16.2% | 21.4% | 18.8% | 29.9% |
| Parvovirus B19 | 1.9% | 9.3% | 15.4% | 21.7% | 19.9% | 31.8% |
| VZV | 2.6% | 11.3% | 16.6% | 21.7% | 18.2% | 29.8% |

The "serological survey" structure refers to the structure of the pathogen-specific measures available. VZV: varicella-zoster virus.

### MSIRW model with boosting and age-specific waning (MSIRWb-ext AW)

In Goeyvaerts *et al*. (2011), different compartmental models were considered to infer on basic immunological processes for parvovirus B19 (Goeyvaerts et al. 2011). In the MSIRWb-ext AW model, they considered waning of disease-acquired antibodies. Individuals move at a rate $\varepsilon(a)$ from a high immunity state R to a low immunity state W in which they are still protected from infection however categorized as being seronegative, that is, with antibody levels falling below the serostatus threshold (see Figure S2). In addition, they assumed that low immunity can be boosted by exposure to infectious individuals. The boosting rate and the force of infection are proportional with a proportionality constant $\varphi$, such that the rate at which individuals move back from W to R equals $\varphi \cdot \lambda(a)$. By solving the corresponding set of differential equations, one finds that the fraction in state S equals:

$$s(a) = \exp\left(-\int_A^a \lambda(u)\,\mathrm{d}u\right) \text{ if } a > A.$$

Approximating $r(a)$ by $1 - s(a) - w(a)$, $\forall a > A$, assuming $i(a)$ is small relative to $s(a)$ and $w(a)$, the following expression is obtained for the proportion seropositives (if $a > A$):

$$r(a) = \int_A^a \left\{(1-\varphi)\,\lambda(u)\exp\left(-\int_A^u \lambda(v)\mathrm{d}v\right) + \varphi\lambda(u)\right\} \exp\left(-\int_u^a \{\varphi\lambda(v) + \varepsilon(v)\}\,\mathrm{d}v\right)\mathrm{d}u$$

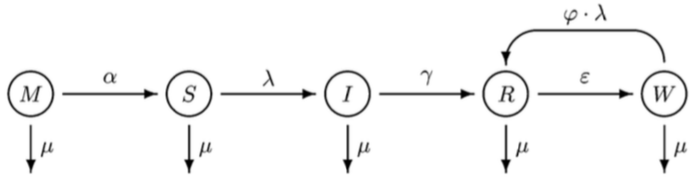

**Figure S2.** Illustration of the MSIRWb-ext compartmental model.

The estimation method assumes endemic and demographic equilibrium. The transmission rates $\beta(a, a')$ are assumed to be directly proportional to age-specific rates of making social contact, $c(a, a')$, with a disease-specific proportionality factor $q$ ("constant proportionality assumption"): $\beta(a, a') = q \cdot c(a, a')$. The contact rates are estimated from the POLYMOD contact survey using a non-parametric model (Goeyvaerts et al. 2010). Since the above integral equation has no closed-form solution, we turn to discrete age classes to estimate the parameters. Through an iterative procedure, the Bernoulli log-likelihood for the serological data is maximized. The waning rate is modelled as a piecewise constant function with a cut-off point at predetermined age $H$: $\varepsilon(a) = \varepsilon_1$, if $a \in (A, H)$, and $\varepsilon(a) = \varepsilon_2$, if $a \geq H$. Comparing the overall likelihood for different values of $H$ (from 5 to 50 years in 5 years steps) led to the choice of $H = 35$ years.

### Calculations of the key epidemiological parameters

- Calculation of the overall prevalence: $\hat{p}_{overall} = \frac{\sum_a \hat{p}_a \times f_a}{\sum_a f_a}$ where $\hat{p}_a$ is the estimated age-specific prevalence and $f_a$ the proportion of individuals aged $a$ from the population distribution.
- Calculation of the overall force of infection: $\hat{\lambda}_{overall} = \frac{\sum_a \hat{\lambda}_a \times f_a \times (1-\hat{p}_a)}{\sum_a f_a \times (1-\hat{p}_a)}$ where $\hat{\lambda}_a$ is the estimated age-specific force of infection, $\hat{p}_a$ is the estimated age-specific prevalence and $f_a$ the proportion of individuals aged $a$ from the population distribution.
- Calculation of $R_0$:
  - Exponentially damped model: $1 + \frac{L}{A_{inf}}$
  - MSIRWb-ext AW model (Goeyvaerts et al. 2011): dominant eigenvalue of $G = \frac{ND}{L}\left(a_{[i+1]} - a_{[i]}\right)\beta_{ij}$ where $\beta_{ij}$ denotes the per capita rate at which an individual of age class $j$ makes an effective contact with a person of age class $i$ per year.
- Calculation of the mean age at infection:
  - Exponentially damped model: $A_{inf} = \int_0^U [1 - \pi(a)]da + [1 - \pi(U)] \times (L - U)$ in case that the data are observed up to a certain age $U$ $(U \leq L)$
  - MSIRWb-ext AW model (Goeyvaerts et al. 2011): $A_{inf} = \frac{\int_0^\infty a\lambda(a)s(a)N(a)/N(0)da}{n_{SI}}$ with $n_{SI} = \int_0^\infty \lambda(a)s(a)N(a)/N(0)da$ the average number of infections per person during their lifetime
- Calculation of $R_{eff}$ (MSIRWb-ext AW model, (Santermans et al. 2015)): dominant eigenvalue of $G \times s(a)$

### Estimates of key epidemiological parameters

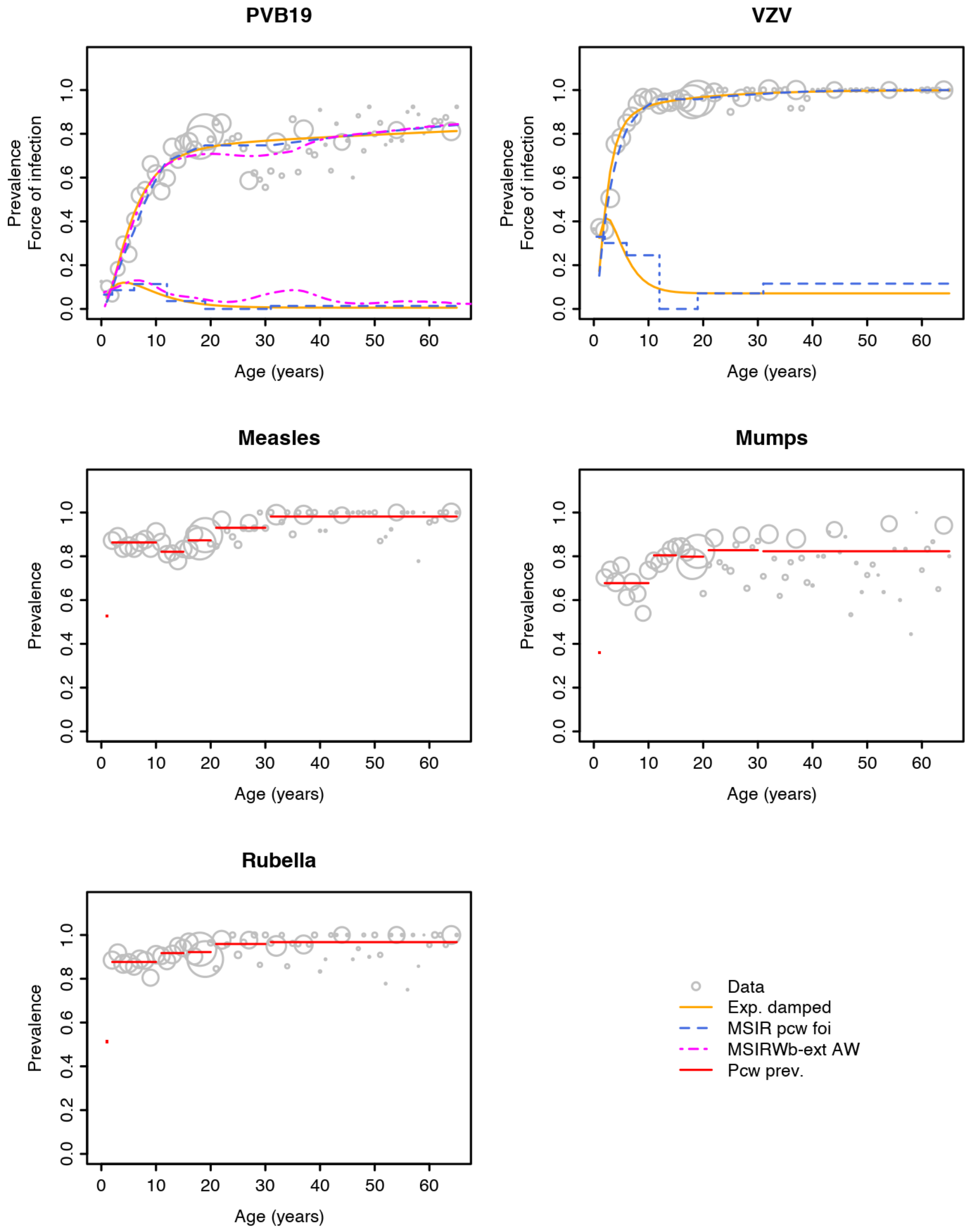

**Figure S3.** Estimated prevalence and force of infection for each of the pathogens.

Points are the observed data with circles proportional to sample size. PVB19: parvovirus B19; VZV: varicella-zoster virus; Exp. damped: exponentially damped model; MSIR - pwc foi: MSIR model with piecewise constant force of infection; MSIRWb-ext AW: MSIR model with boosting and age-specific waning; Pcw. prev: logistic model with piecewise constant prevalence.

**Table S2.** Estimates of the overall prevalence and prevalence by age groups by model and infection

| | Prevalence | | | | | | |
|---|---|---|---|---|---|---|---|
| | [1,2) yr | [2,6) yrs | [6,12) yrs | [12,19) yrs | [19,31) yrs | [31,65] yrs | Overall |
| **MSIR with piecewise constant force of infection** | | | | | | | |
| VZV | 18.3% | 64.8% | 91.0% | 95.7% | 97.1% | 99.5% | 95.0% |
| B19 | 3.8% | 23.3% | 53.8% | 70.9% | 74.8% | 79.5% | 71.5% |
| **Exponentially damped model** | | | | | | | |
| VZV | 17.3% | 65.7% | 90.6% | 95.3% | 97.6% | 99.4% | 95.0% |
| B19 | 3.3% | 24.1% | 55.4% | 69.9% | 75.5% | 79.1% | 71.5% |
| **MSIRWb-ext AW** B19 | 4.1% | 24.2% | 56.7% | 70.1% | 70.4% | 78.6% | 70.5% |
| **Logistic model** | | | | | | | |
| Measles | 52.7% | 86.3% | 85.6% | 84.3% | 92.1% | 98.2% | 93.3% |
| Mumps | 36.0% | 67.7% | 70.0% | 80.2% | 82.3% | 82.3% | 79.6% |
| Rubella | 51.2% | 87.7% | 88.4% | 91.9% | 95.3% | 96.7% | 94.2% |

All models are applied to data with integer age values.

**Table S3.** Estimates of the overall force of infection and force of infection by age groups by model and infection

| | Force of infection | | | | | | |
|---|---|---|---|---|---|---|---|
| | [1,2) yr | [2,6) yrs | [6,12) yrs | [12,19) yrs | [19,31) yrs | [31,65] yrs | Overall |
| **MSIR with piecewise constant force of infection** | | | | | | | |
| VZV | 0.404 | 0.337 | 0.200 | 0.000 | 0.076 | 0.113 | 0.262 |
| B19 | 0.077 | 0.104 | 0.100 | 0.035 | 0.000 | 0.014 | 0.042 |
| **Exp. damped model** | | | | | | | |
| VZV | 0.325 | 0.385 | 0.168 | 0.081 | 0.071 | 0.071 | 0.260 |
| B19 | 0.061 | 0.111 | 0.091 | 0.039 | 0.012 | 0.006 | 0.040 |
| **MSIRWb-ext AW** B19 | 0.090 | 0.114 | 0.124 | 0.060 | 0.045 | 0.052 | 0.071 |

All models are applied to data with integer age values.

**Table S4.** Estimates of $R_0$, average age at infection, and $R_{eff}$, by model and infection

| | $R_0$ | Average age at infection | $R_{eff}$ |
|---|---|---|---|
| **Exp. damped model** | | | |
| VZV | 22.92 | 3.6 | - |
| B19 | 4.65 | 21.6 | - |
| **MSIRWb-ext AW** B19 | 3.94 | 10.7 | 1.06 |

All models are applied to data with integer age values.

### Precision values

In Tables S5-12, the following abbreviations are used:

survey: age structure derived from the serological survey data; uniform: uniform age structure; population: age structure of the Belgian population; OverallPrev: overall seroprevalence; Prev1 to Prev6: prevalence by age group ([1,2), [2,6), [6,12), [12,19), [19,31), and [31,65] years); OverallFoi: overall force of infection; Foi1 to Foi6: force of infection by age group ([1,2), [2,6), [6,12), [12,19), [19,31), and [31,65] years); $R_0$: basic reproduction number; $R_{eff}$: effective reproduction number; ASI: average age at infection.

**Table S5.** Measles serological data – Logistic model with piecewise constant prevalence

| Size | Structure | OverallPrev | Prev1 | Prev2 | Prev3 | Prev4 | Prev5 | Prev6 |
|---|---|---|---|---|---|---|---|---|
| 1650 | survey | 0.0106 | 0.1579 | 0.0330 | 0.0292 | 0.0372 | 0.0289 | 0.0119 |
| 3300 | survey | 0.0082 | 0.1104 | 0.0214 | 0.0193 | 0.0243 | 0.0214 | 0.0084 |
| 6600 | survey | 0.0054 | 0.0796 | 0.0160 | 0.0143 | 0.0166 | 0.0153 | 0.0053 |
| 9900 | survey | 0.0041 | 0.0628 | 0.0136 | 0.0115 | 0.0142 | 0.0125 | 0.0047 |
| 13200 | survey | 0.0038 | 0.0552 | 0.0111 | 0.0100 | 0.0125 | 0.0110 | 0.0044 |
| 19800 | survey | 0.0031 | 0.0478 | 0.0096 | 0.0081 | 0.0091 | 0.0085 | 0.0033 |
| 1650 | uniform | 0.0110 | 0.2200 | 0.0434 | 0.0395 | 0.0463 | 0.0270 | 0.0083 |
| 3300 | uniform | 0.0084 | 0.1353 | 0.0300 | 0.0274 | 0.0335 | 0.0197 | 0.0066 |
| 6600 | uniform | 0.0055 | 0.0990 | 0.0215 | 0.0180 | 0.0226 | 0.0145 | 0.0042 |
| 9900 | uniform | 0.0047 | 0.0789 | 0.0174 | 0.0153 | 0.0193 | 0.0118 | 0.0032 |
| 13200 | uniform | 0.0038 | 0.0665 | 0.0153 | 0.0134 | 0.0169 | 0.0099 | 0.0032 |
| 19800 | uniform | 0.0032 | 0.0559 | 0.0130 | 0.0110 | 0.0134 | 0.0084 | 0.0027 |
| 1650 | population | 0.0109 | 0.2045 | 0.0447 | 0.0425 | 0.0468 | 0.0273 | 0.0086 |
| 3300 | population | 0.0084 | 0.1423 | 0.0336 | 0.0287 | 0.0348 | 0.0196 | 0.0059 |
| 6600 | population | 0.0056 | 0.1023 | 0.0219 | 0.0190 | 0.0234 | 0.0147 | 0.0042 |
| 9900 | population | 0.0048 | 0.0833 | 0.0187 | 0.0160 | 0.0198 | 0.0119 | 0.0032 |
| 13200 | population | 0.0039 | 0.0710 | 0.0161 | 0.0144 | 0.0156 | 0.0102 | 0.0031 |
| 19800 | population | 0.0031 | 0.0625 | 0.0141 | 0.0116 | 0.0146 | 0.0081 | 0.0025 |

**Table S6.** Mumps serological data – Logistic model with piecewise constant prevalence

| Size | Structure | OverallPrev | Prev1 | Prev2 | Prev3 | Prev4 | Prev5 | Prev6 |
|---|---|---|---|---|---|---|---|---|
| 1650 | survey | 0.0207 | 0.1463 | 0.0439 | 0.0364 | 0.0367 | 0.0442 | 0.0316 |
| 3300 | survey | 0.0153 | 0.0964 | 0.0320 | 0.0261 | 0.0251 | 0.0304 | 0.0233 |
| 6600 | survey | 0.0112 | 0.0693 | 0.0212 | 0.0182 | 0.0171 | 0.0232 | 0.0164 |
| 9900 | survey | 0.0093 | 0.0615 | 0.0184 | 0.0160 | 0.0149 | 0.0176 | 0.0138 |
| 13200 | survey | 0.0075 | 0.0537 | 0.0161 | 0.0138 | 0.0129 | 0.0167 | 0.0111 |
| 19800 | survey | 0.0065 | 0.0408 | 0.0133 | 0.0110 | 0.0099 | 0.0132 | 0.0104 |
| 1650 | uniform | 0.0187 | 0.1800 | 0.0589 | 0.0527 | 0.0515 | 0.0443 | 0.0240 |
| 3300 | uniform | 0.0136 | 0.1300 | 0.0406 | 0.0350 | 0.0371 | 0.0284 | 0.0181 |
| 6600 | uniform | 0.0092 | 0.0941 | 0.0303 | 0.0248 | 0.0243 | 0.0211 | 0.0132 |
| 9900 | uniform | 0.0078 | 0.0724 | 0.0243 | 0.0198 | 0.0210 | 0.0177 | 0.0106 |
| 13200 | uniform | 0.0065 | 0.0640 | 0.0194 | 0.0169 | 0.0178 | 0.0157 | 0.0083 |
| 19800 | uniform | 0.0054 | 0.0551 | 0.0181 | 0.0154 | 0.0156 | 0.0121 | 0.0076 |
| 1650 | population | 0.0182 | 0.2045 | 0.0617 | 0.0538 | 0.0508 | 0.0427 | 0.0226 |
| 3300 | population | 0.0140 | 0.1364 | 0.0439 | 0.0367 | 0.0358 | 0.0278 | 0.0171 |
| 6600 | population | 0.0093 | 0.0996 | 0.0313 | 0.0260 | 0.0263 | 0.0211 | 0.0125 |
| 9900 | population | 0.0078 | 0.0758 | 0.0244 | 0.0205 | 0.0215 | 0.0168 | 0.0107 |
| 13200 | population | 0.0063 | 0.0668 | 0.0211 | 0.0181 | 0.0171 | 0.0150 | 0.0081 |
| 19800 | population | 0.0054 | 0.0606 | 0.0189 | 0.0158 | 0.0149 | 0.0117 | 0.0075 |

**Table S7.** Rubella serological data – Logistic model with piecewise constant prevalence

| Size | Structure | OverallPrev | Prev1 | Prev2 | Prev3 | Prev4 | Prev5 | Prev6 |
|---|---|---|---|---|---|---|---|---|
| 1650 | survey | 0.0111 | 0.1548 | 0.0336 | 0.0279 | 0.0251 | 0.0220 | 0.0152 |
| 3300 | survey | 0.0085 | 0.1084 | 0.0215 | 0.0183 | 0.0174 | 0.0171 | 0.0120 |
| 6600 | survey | 0.0057 | 0.0723 | 0.0161 | 0.0139 | 0.0121 | 0.0123 | 0.0076 |
| 9900 | survey | 0.0046 | 0.0622 | 0.0133 | 0.0112 | 0.0110 | 0.0100 | 0.0061 |
| 13200 | survey | 0.0037 | 0.0557 | 0.0113 | 0.0095 | 0.0090 | 0.0083 | 0.0054 |
| 19800 | survey | 0.0033 | 0.0462 | 0.0091 | 0.0078 | 0.0072 | 0.0073 | 0.0043 |
| 1650 | uniform | 0.0105 | 0.2000 | 0.0423 | 0.0370 | 0.0345 | 0.0215 | 0.0120 |
| 3300 | uniform | 0.0074 | 0.1300 | 0.0312 | 0.0256 | 0.0240 | 0.0154 | 0.0080 |
| 6600 | uniform | 0.0055 | 0.0967 | 0.0204 | 0.0177 | 0.0168 | 0.0118 | 0.0057 |
| 9900 | uniform | 0.0046 | 0.0822 | 0.0170 | 0.0146 | 0.0146 | 0.0088 | 0.0043 |
| 13200 | uniform | 0.0037 | 0.0678 | 0.0151 | 0.0131 | 0.0117 | 0.0075 | 0.0038 |
| 19800 | uniform | 0.0032 | 0.0593 | 0.0122 | 0.0103 | 0.0100 | 0.0062 | 0.0034 |
| 1650 | population | 0.0105 | 0.2045 | 0.0411 | 0.0376 | 0.0346 | 0.0231 | 0.0113 |
| 3300 | population | 0.0075 | 0.1477 | 0.0329 | 0.0269 | 0.0241 | 0.0152 | 0.0077 |
| 6600 | population | 0.0053 | 0.1026 | 0.0213 | 0.0184 | 0.0175 | 0.0114 | 0.0058 |
| 9900 | population | 0.0045 | 0.0871 | 0.0177 | 0.0146 | 0.0146 | 0.0089 | 0.0043 |
| 13200 | population | 0.0037 | 0.0739 | 0.0158 | 0.0136 | 0.0122 | 0.0076 | 0.0036 |
| 19800 | population | 0.0033 | 0.0625 | 0.0137 | 0.0113 | 0.0106 | 0.0065 | 0.0033 |

**Table S8.** VZV serological data – Exponentially damped model

| Size | Structure | OverallPrev | Prev1 | Prev2 | Prev3 | Prev4 | Prev5 | Prev6 |
|---|---|---|---|---|---|---|---|---|
| 1650 | survey | 0.0071 | 0.0520 | 0.0591 | 0.0250 | 0.0142 | 0.0102 | 0.0056 |
| 3300 | survey | 0.0049 | 0.0333 | 0.0386 | 0.0160 | 0.0103 | 0.0069 | 0.0043 |
| 6600 | survey | 0.0037 | 0.0240 | 0.0285 | 0.0126 | 0.0075 | 0.0048 | 0.0028 |
| 9900 | survey | 0.0026 | 0.0220 | 0.0242 | 0.0103 | 0.0059 | 0.0031 | 0.0018 |
| 13200 | survey | 0.0023 | 0.0184 | 0.0206 | 0.0087 | 0.0052 | 0.0032 | 0.0020 |
| 19800 | survey | 0.0021 | 0.0142 | 0.0156 | 0.0068 | 0.0045 | 0.0027 | 0.0016 |
| 1650 | uniform | 0.0085 | 0.0716 | 0.0785 | 0.0320 | 0.0182 | 0.0091 | 0.0041 |
| 3300 | uniform | 0.0060 | 0.0493 | 0.0585 | 0.0212 | 0.0131 | 0.0067 | 0.0032 |
| 6600 | uniform | 0.0040 | 0.0328 | 0.0386 | 0.0152 | 0.0095 | 0.0047 | 0.0021 |
| 9900 | uniform | 0.0038 | 0.0268 | 0.0307 | 0.0127 | 0.0077 | 0.0038 | 0.0018 |
| 13200 | uniform | 0.0031 | 0.0234 | 0.0280 | 0.0114 | 0.0064 | 0.0032 | 0.0015 |
| 19800 | uniform | 0.0023 | 0.0195 | 0.0227 | 0.0084 | 0.0056 | 0.0026 | 0.0013 |
| 1650 | population | 0.0090 | 0.0779 | 0.0881 | 0.0323 | 0.0187 | 0.0091 | 0.0041 |
| 3300 | population | 0.0057 | 0.0508 | 0.0572 | 0.0226 | 0.0131 | 0.0065 | 0.0029 |
| 6600 | population | 0.0042 | 0.0352 | 0.0412 | 0.0154 | 0.0101 | 0.0047 | 0.0022 |
| 9900 | population | 0.0035 | 0.0284 | 0.0325 | 0.0128 | 0.0082 | 0.0039 | 0.0017 |
| 13200 | population | 0.0029 | 0.0255 | 0.0282 | 0.0115 | 0.0069 | 0.0032 | 0.0014 |
| 19800 | population | 0.0025 | 0.0205 | 0.0252 | 0.0091 | 0.0056 | 0.0027 | 0.0012 |

| Size | Structure | OverallFoi | Foi1 | Foi2 | Foi3 | Foi4 | Foi5 | Foi6 |
|---|---|---|---|---|---|---|---|---|
| 1650 | survey | 0.0353 | 0.0906 | 0.0518 | 0.0473 | 0.0359 | 0.0434 | 0.0441 |
| 3300 | survey | 0.0239 | 0.0603 | 0.0326 | 0.0350 | 0.0234 | 0.0288 | 0.0291 |
| 6600 | survey | 0.0179 | 0.0434 | 0.0236 | 0.0244 | 0.0146 | 0.0187 | 0.0189 |
| 9900 | survey | 0.0129 | 0.0395 | 0.0185 | 0.0230 | 0.0101 | 0.0138 | 0.0141 |
| 13200 | survey | 0.0114 | 0.0325 | 0.0171 | 0.0182 | 0.0115 | 0.0146 | 0.0148 |
| 19800 | survey | 0.0100 | 0.0252 | 0.0120 | 0.0153 | 0.0082 | 0.0108 | 0.0109 |
| 1650 | uniform | 0.0416 | 0.1268 | 0.0658 | 0.0616 | 0.0280 | 0.0330 | 0.0338 |
| 3300 | uniform | 0.0294 | 0.0891 | 0.0439 | 0.0470 | 0.0192 | 0.0245 | 0.0251 |
| 6600 | uniform | 0.0195 | 0.0601 | 0.0299 | 0.0341 | 0.0134 | 0.0163 | 0.0168 |
| 9900 | uniform | 0.0176 | 0.0474 | 0.0257 | 0.0268 | 0.0102 | 0.0133 | 0.0137 |
| 13200 | uniform | 0.0150 | 0.0425 | 0.0220 | 0.0227 | 0.0086 | 0.0115 | 0.0118 |
| 19800 | uniform | 0.0113 | 0.0352 | 0.0174 | 0.0198 | 0.0070 | 0.0089 | 0.0092 |
| 1650 | population | 0.0433 | 0.1410 | 0.0689 | 0.0659 | 0.0273 | 0.0339 | 0.0351 |
| 3300 | population | 0.0279 | 0.0907 | 0.0452 | 0.0487 | 0.0184 | 0.0251 | 0.0257 |
| 6600 | population | 0.0201 | 0.0629 | 0.0309 | 0.0362 | 0.0133 | 0.0162 | 0.0166 |
| 9900 | population | 0.0172 | 0.0509 | 0.0249 | 0.0282 | 0.0098 | 0.0132 | 0.0135 |
| 13200 | population | 0.0143 | 0.0453 | 0.0241 | 0.0253 | 0.0086 | 0.0113 | 0.0115 |
| 19800 | population | 0.0118 | 0.0371 | 0.0186 | 0.0211 | 0.0072 | 0.0090 | 0.0092 |

| Size | Structure | $R_0$ | ASI |
|---|---|---|---|
| 1650 | survey | 3.0545 | 0.4995 |
| 3300 | survey | 2.1016 | 0.3472 |
| 6600 | survey | 1.5552 | 0.2533 |
| 9900 | survey | 1.1384 | 0.1840 |
| 13200 | survey | 0.9811 | 0.1599 |
| 19800 | survey | 0.8942 | 0.1464 |
| 1650 | uniform | 3.7604 | 0.5920 |
| 3300 | uniform | 2.5140 | 0.4081 |
| 6600 | uniform | 1.7357 | 0.2816 |
| 9900 | uniform | 1.6336 | 0.2674 |
| 13200 | uniform | 1.3314 | 0.2190 |
| 19800 | uniform | 1.0348 | 0.1679 |
| 1650 | population | 3.9463 | 0.6310 |
| 3300 | population | 2.4735 | 0.4046 |
| 6600 | population | 1.8329 | 0.3006 |
| 9900 | population | 1.5411 | 0.2524 |
| 13200 | population | 1.2795 | 0.2096 |
| 19800 | population | 1.0671 | 0.1741 |

**Table S9.** VZV serological data – MSIR piecewise constant force of infection

| Size | Structure | OverallPrev | Prev1 | Prev2 | Prev3 | Prev4 | Prev5 | Prev6 |
|---|---|---|---|---|---|---|---|---|
| 1650 | survey | 0.0073 | 0.0575 | 0.0610 | 0.0251 | 0.0162 | 0.0136 | 0.0046 |
| 3300 | survey | 0.0049 | 0.0359 | 0.0402 | 0.0177 | 0.0115 | 0.0101 | 0.0038 |
| 6600 | survey | 0.0038 | 0.0279 | 0.0292 | 0.0128 | 0.0079 | 0.0071 | 0.0028 |
| 9900 | survey | 0.0030 | 0.0221 | 0.0236 | 0.0103 | 0.0068 | 0.0054 | 0.0023 |
| 13200 | survey | 0.0024 | 0.0210 | 0.0204 | 0.0090 | 0.0057 | 0.0047 | 0.0020 |
| 19800 | survey | 0.0020 | 0.0158 | 0.0159 | 0.0073 | 0.0050 | 0.0040 | 0.0016 |
| 1650 | uniform | 0.0087 | 0.0798 | 0.0764 | 0.0352 | 0.0226 | 0.0137 | 0.0038 |
| 3300 | uniform | 0.0060 | 0.0543 | 0.0557 | 0.0254 | 0.0158 | 0.0102 | 0.0029 |
| 6600 | uniform | 0.0042 | 0.0342 | 0.0387 | 0.0172 | 0.0111 | 0.0072 | 0.0022 |
| 9900 | uniform | 0.0036 | 0.0314 | 0.0310 | 0.0137 | 0.0097 | 0.0055 | 0.0017 |
| 13200 | uniform | 0.0032 | 0.0259 | 0.0287 | 0.0119 | 0.0081 | 0.0050 | 0.0014 |
| 19800 | uniform | 0.0023 | 0.0217 | 0.0221 | 0.0097 | 0.0062 | 0.0038 | 0.0012 |
| 1650 | population | 0.0089 | 0.0842 | 0.0880 | 0.0360 | 0.0222 | 0.0140 | 0.0037 |
| 3300 | population | 0.0061 | 0.0564 | 0.0587 | 0.0249 | 0.0167 | 0.0101 | 0.0028 |
| 6600 | population | 0.0043 | 0.0373 | 0.0417 | 0.0173 | 0.0116 | 0.0071 | 0.0021 |
| 9900 | population | 0.0036 | 0.0305 | 0.0339 | 0.0142 | 0.0098 | 0.0056 | 0.0016 |
| 13200 | population | 0.0031 | 0.0292 | 0.0305 | 0.0131 | 0.0080 | 0.0049 | 0.0014 |
| 19800 | population | 0.0025 | 0.0219 | 0.0241 | 0.0104 | 0.0065 | 0.0039 | 0.0012 |

| Size | Structure | OverallFoi | Foi1 | Foi2 | Foi3 | Foi4 | Foi5 | Foi6 |
|---|---|---|---|---|---|---|---|---|
| 1650 | survey | 0.0383 | 0.1416 | 0.1307 | 0.1137 | 0.0711 | 0.0894 | 0.3651 |
| 3300 | survey | 0.0263 | 0.0879 | 0.0910 | 0.0853 | 0.0461 | 0.0774 | 0.2816 |
| 6600 | survey | 0.0200 | 0.0683 | 0.0560 | 0.0573 | 0.0389 | 0.0570 | 0.1015 |
| 9900 | survey | 0.0152 | 0.0542 | 0.0502 | 0.0470 | 0.0285 | 0.0482 | 0.0778 |
| 13200 | survey | 0.0134 | 0.0514 | 0.0439 | 0.0417 | 0.0253 | 0.0379 | 0.0610 |
| 19800 | survey | 0.0110 | 0.0386 | 0.0365 | 0.0316 | 0.0207 | 0.0294 | 0.0480 |
| 1650 | uniform | 0.0461 | 0.1983 | 0.1555 | 0.1422 | 0.0919 | 0.0892 | 0.2131 |
| 3300 | uniform | 0.0340 | 0.1339 | 0.1216 | 0.1100 | 0.0661 | 0.0717 | 0.1013 |
| 6600 | uniform | 0.0240 | 0.0837 | 0.0834 | 0.0807 | 0.0454 | 0.0584 | 0.0679 |
| 9900 | uniform | 0.0189 | 0.0772 | 0.0660 | 0.0626 | 0.0420 | 0.0463 | 0.0496 |
| 13200 | uniform | 0.0159 | 0.0633 | 0.0516 | 0.0536 | 0.0361 | 0.0402 | 0.0430 |
| 19800 | uniform | 0.0125 | 0.0531 | 0.0471 | 0.0430 | 0.0266 | 0.0317 | 0.0364 |
| 1650 | population | 0.0453 | 0.2088 | 0.1763 | 0.1616 | 0.0937 | 0.0819 | 0.2257 |
| 3300 | population | 0.0336 | 0.1386 | 0.1282 | 0.1117 | 0.0661 | 0.0671 | 0.0989 |
| 6600 | population | 0.0233 | 0.0915 | 0.0869 | 0.0866 | 0.0446 | 0.0593 | 0.0645 |
| 9900 | population | 0.0190 | 0.0748 | 0.0691 | 0.0652 | 0.0405 | 0.0445 | 0.0540 |
| 13200 | population | 0.0158 | 0.0715 | 0.0587 | 0.0567 | 0.0379 | 0.0383 | 0.0439 |
| 19800 | population | 0.0131 | 0.0537 | 0.0486 | 0.0471 | 0.0255 | 0.0315 | 0.0331 |

**Table S10.** Parvovirus B19 serological data – Exponentially damped model

| Size | Structure | OverallPrev | Prev1 | Prev2 | Prev3 | Prev4 | Prev5 | Prev6 |
|---|---|---|---|---|---|---|---|---|
| 1650 | survey | 0.0215 | 0.0094 | 0.0415 | 0.0391 | 0.0305 | 0.0285 | 0.0294 |
| 3300 | survey | 0.0148 | 0.0074 | 0.0321 | 0.0298 | 0.0213 | 0.0202 | 0.0208 |
| 6600 | survey | 0.0106 | 0.0049 | 0.0213 | 0.0203 | 0.0156 | 0.0148 | 0.0148 |
| 9900 | survey | 0.0092 | 0.0039 | 0.0175 | 0.0163 | 0.0115 | 0.0120 | 0.0138 |
| 13200 | survey | 0.0073 | 0.0033 | 0.0144 | 0.0135 | 0.0099 | 0.0105 | 0.0109 |
| 19800 | survey | 0.0071 | 0.0029 | 0.0132 | 0.0117 | 0.0087 | 0.0085 | 0.0100 |
| 1650 | uniform | 0.0199 | 0.0126 | 0.0552 | 0.0511 | 0.0376 | 0.0321 | 0.0226 |
| 3300 | uniform | 0.0152 | 0.0086 | 0.0392 | 0.0407 | 0.0268 | 0.0218 | 0.0166 |
| 6600 | uniform | 0.0097 | 0.0062 | 0.0278 | 0.0281 | 0.0175 | 0.0160 | 0.0120 |
| 9900 | uniform | 0.0082 | 0.0048 | 0.0219 | 0.0203 | 0.0150 | 0.0135 | 0.0098 |
| 13200 | uniform | 0.0069 | 0.0042 | 0.0190 | 0.0184 | 0.0120 | 0.0111 | 0.0079 |
| 19800 | uniform | 0.0055 | 0.0033 | 0.0149 | 0.0145 | 0.0094 | 0.0092 | 0.0074 |
| 1650 | population | 0.0192 | 0.0135 | 0.0570 | 0.0536 | 0.0366 | 0.0300 | 0.0215 |
| 3300 | population | 0.0142 | 0.0090 | 0.0402 | 0.0416 | 0.0277 | 0.0223 | 0.0173 |
| 6600 | population | 0.0097 | 0.0065 | 0.0292 | 0.0269 | 0.0176 | 0.0159 | 0.0117 |
| 9900 | population | 0.0079 | 0.0050 | 0.0233 | 0.0224 | 0.0152 | 0.0130 | 0.0101 |
| 13200 | population | 0.0071 | 0.0043 | 0.0201 | 0.0200 | 0.0128 | 0.0115 | 0.0081 |
| 19800 | population | 0.0059 | 0.0038 | 0.0174 | 0.0162 | 0.0096 | 0.0091 | 0.0076 |

| Size | Structure | OverallFoi | Foi1 | Foi2 | Foi3 | Foi4 | Foi5 | Foi6 |
|---|---|---|---|---|---|---|---|---|
| 1650 | survey | 0.0050 | 0.0167 | 0.0174 | 0.0117 | 0.0117 | 0.0054 | 0.0064 |
| 3300 | survey | 0.0036 | 0.0131 | 0.0130 | 0.0084 | 0.0091 | 0.0040 | 0.0056 |
| 6600 | survey | 0.0025 | 0.0087 | 0.0089 | 0.0057 | 0.0065 | 0.0028 | 0.0038 |
| 9900 | survey | 0.0023 | 0.0070 | 0.0071 | 0.0049 | 0.0053 | 0.0024 | 0.0036 |
| 13200 | survey | 0.0020 | 0.0058 | 0.0059 | 0.0044 | 0.0045 | 0.0020 | 0.0031 |
| 19800 | survey | 0.0018 | 0.0053 | 0.0054 | 0.0034 | 0.0039 | 0.0017 | 0.0025 |
| 1650 | uniform | 0.0037 | 0.0223 | 0.0223 | 0.0131 | 0.0142 | 0.0055 | 0.0065 |
| 3300 | uniform | 0.0030 | 0.0154 | 0.0168 | 0.0093 | 0.0098 | 0.0039 | 0.0051 |
| 6600 | uniform | 0.0019 | 0.0110 | 0.0117 | 0.0065 | 0.0072 | 0.0027 | 0.0038 |
| 9900 | uniform | 0.0018 | 0.0086 | 0.0090 | 0.0055 | 0.0062 | 0.0022 | 0.0033 |
| 13200 | uniform | 0.0014 | 0.0075 | 0.0079 | 0.0045 | 0.0052 | 0.0019 | 0.0027 |
| 19800 | uniform | 0.0013 | 0.0059 | 0.0062 | 0.0036 | 0.0041 | 0.0016 | 0.0021 |
| 1650 | population | 0.0038 | 0.0241 | 0.0240 | 0.0141 | 0.0138 | 0.0055 | 0.0067 |
| 3300 | population | 0.0029 | 0.0161 | 0.0174 | 0.0094 | 0.0101 | 0.0040 | 0.0054 |
| 6600 | population | 0.0021 | 0.0118 | 0.0120 | 0.0069 | 0.0077 | 0.0027 | 0.0040 |
| 9900 | population | 0.0019 | 0.0090 | 0.0097 | 0.0055 | 0.0060 | 0.0022 | 0.0034 |
| 13200 | population | 0.0015 | 0.0077 | 0.0083 | 0.0045 | 0.0053 | 0.0020 | 0.0029 |
| 19800 | population | 0.0013 | 0.0068 | 0.0071 | 0.0036 | 0.0045 | 0.0017 | 0.0023 |

| **Size** | **Structure** | $\mathbf{R_0}$ | **ASI** |
|---|---|---|---|
| 1650 | survey | 0.3125 | 1.8265 |
| 3300 | survey | 0.2283 | 1.3602 |
| 6600 | survey | 0.1537 | 0.9046 |
| 9900 | survey | 0.1415 | 0.8382 |
| 13200 | survey | 0.1198 | 0.7077 |
| 19800 | survey | 0.1081 | 0.6394 |
| 1650 | uniform | 0.2661 | 1.5597 |
| 3300 | uniform | 0.2004 | 1.1785 |
| 6600 | uniform | 0.1320 | 0.7797 |
| 9900 | uniform | 0.1140 | 0.6742 |
| 13200 | uniform | 0.0927 | 0.5481 |
| 19800 | uniform | 0.0817 | 0.4841 |
| 1650 | population | 0.2550 | 1.4957 |
| 3300 | population | 0.1932 | 1.1408 |
| 6600 | population | 0.1361 | 0.8059 |
| 9900 | population | 0.1161 | 0.6863 |
| 13200 | population | 0.0951 | 0.5629 |
| 19800 | population | 0.0843 | 0.4997 |

**Table S11.** Parvovirus B19 serological data – MSIR piecewise constant force of infection

| Size | Structure | OverallPrev | Prev1 | Prev2 | Prev3 | Prev4 | Prev5 | Prev6 |
|---|---|---|---|---|---|---|---|---|
| 1650 | survey | 0.0205 | 0.0277 | 0.0483 | 0.0498 | 0.0378 | 0.0341 | 0.0284 |
| 3300 | survey | 0.0150 | 0.0203 | 0.0383 | 0.0350 | 0.0246 | 0.0235 | 0.0199 |
| 6600 | survey | 0.0102 | 0.0149 | 0.0264 | 0.0241 | 0.0195 | 0.0178 | 0.0147 |
| 9900 | survey | 0.0092 | 0.0118 | 0.0204 | 0.0199 | 0.0153 | 0.0140 | 0.0130 |
| 13200 | survey | 0.0074 | 0.0099 | 0.0183 | 0.0169 | 0.0130 | 0.0118 | 0.0105 |
| 19800 | survey | 0.0074 | 0.0080 | 0.0156 | 0.0142 | 0.0099 | 0.0098 | 0.0099 |
| 1650 | uniform | 0.0199 | 0.0331 | 0.0633 | 0.0569 | 0.0458 | 0.0371 | 0.0225 |
| 3300 | uniform | 0.0144 | 0.0232 | 0.0470 | 0.0471 | 0.0370 | 0.0272 | 0.0161 |
| 6600 | uniform | 0.0100 | 0.0163 | 0.0322 | 0.0304 | 0.0238 | 0.0183 | 0.0128 |
| 9900 | uniform | 0.0079 | 0.0140 | 0.0261 | 0.0248 | 0.0200 | 0.0155 | 0.0099 |
| 13200 | uniform | 0.0067 | 0.0121 | 0.0235 | 0.0225 | 0.0183 | 0.0131 | 0.0081 |
| 19800 | uniform | 0.0057 | 0.0095 | 0.0187 | 0.0183 | 0.0138 | 0.0103 | 0.0075 |
| 1650 | population | 0.0196 | 0.0387 | 0.0683 | 0.0614 | 0.0473 | 0.0356 | 0.0223 |
| 3300 | population | 0.0141 | 0.0253 | 0.0473 | 0.0468 | 0.0385 | 0.0266 | 0.0159 |
| 6600 | population | 0.0097 | 0.0181 | 0.0343 | 0.0330 | 0.0245 | 0.0181 | 0.0120 |
| 9900 | population | 0.0079 | 0.0148 | 0.0272 | 0.0256 | 0.0196 | 0.0150 | 0.0098 |
| 13200 | population | 0.0072 | 0.0128 | 0.0241 | 0.0235 | 0.0189 | 0.0131 | 0.0079 |
| 19800 | population | 0.0058 | 0.0103 | 0.0200 | 0.0202 | 0.0148 | 0.0103 | 0.0074 |

| Size | Structure | OverallFoi | Foi1 | Foi2 | Foi3 | Foi4 | Foi5 | Foi6 |
|---|---|---|---|---|---|---|---|---|
| 1650 | survey | 0.0053 | 0.0578 | 0.0453 | 0.0430 | 0.0355 | 0.0123 | 0.0120 |
| 3300 | survey | 0.0037 | 0.0423 | 0.0315 | 0.0298 | 0.0291 | 0.0098 | 0.0091 |
| 6600 | survey | 0.0027 | 0.0311 | 0.0241 | 0.0240 | 0.0212 | 0.0073 | 0.0057 |
| 9900 | survey | 0.0025 | 0.0244 | 0.0211 | 0.0199 | 0.0173 | 0.0055 | 0.0051 |
| 13200 | survey | 0.0021 | 0.0205 | 0.0160 | 0.0165 | 0.0139 | 0.0047 | 0.0048 |
| 19800 | survey | 0.0018 | 0.0167 | 0.0137 | 0.0144 | 0.0119 | 0.0040 | 0.0036 |
| 1650 | uniform | 0.0047 | 0.0692 | 0.0598 | 0.0575 | 0.0375 | 0.0141 | 0.0109 |
| 3300 | uniform | 0.0036 | 0.0484 | 0.0414 | 0.0457 | 0.0338 | 0.0108 | 0.0075 |
| 6600 | uniform | 0.0023 | 0.0339 | 0.0288 | 0.0300 | 0.0269 | 0.0082 | 0.0054 |
| 9900 | uniform | 0.0018 | 0.0291 | 0.0235 | 0.0244 | 0.0206 | 0.0063 | 0.0044 |
| 13200 | uniform | 0.0016 | 0.0251 | 0.0206 | 0.0216 | 0.0195 | 0.0051 | 0.0034 |
| 19800 | uniform | 0.0014 | 0.0197 | 0.0154 | 0.0173 | 0.0150 | 0.0044 | 0.0031 |
| 1650 | population | 0.0046 | 0.0806 | 0.0641 | 0.0583 | 0.0388 | 0.0138 | 0.0103 |
| 3300 | population | 0.0036 | 0.0527 | 0.0461 | 0.0474 | 0.0339 | 0.0104 | 0.0074 |
| 6600 | population | 0.0024 | 0.0377 | 0.0304 | 0.0297 | 0.0258 | 0.0082 | 0.0054 |
| 9900 | population | 0.0019 | 0.0309 | 0.0256 | 0.0259 | 0.0225 | 0.0065 | 0.0045 |
| 13200 | population | 0.0017 | 0.0266 | 0.0223 | 0.0224 | 0.0185 | 0.0056 | 0.0038 |
| 19800 | population | 0.0015 | 0.0214 | 0.0184 | 0.0189 | 0.0155 | 0.0046 | 0.0032 |

**Table S12.** Parvovirus B19 serological data – MSIRWb-ext AW model

| Size | Structure | OverallPrev | Prev1 | Prev2 | Prev3 | Prev4 | Prev5 | Prev6 |
|---|---|---|---|---|---|---|---|---|
| 1650 | survey | 0.0267 | 0.0078 | 0.0351 | 0.0440 | 0.0311 | 0.0430 | 0.0406 |
| 3300 | survey | 0.0194 | 0.0060 | 0.0273 | 0.0362 | 0.0207 | 0.0351 | 0.0297 |
| 6600 | survey | 0.0140 | 0.0036 | 0.0163 | 0.0219 | 0.0149 | 0.0243 | 0.0209 |
| 9900 | survey | 0.0122 | 0.0032 | 0.0147 | 0.0182 | 0.0124 | 0.0210 | 0.0192 |
| 13200 | survey | 0.0092 | 0.0030 | 0.0133 | 0.0163 | 0.0105 | 0.0203 | 0.0145 |
| 19800 | survey | 0.0083 | 0.0024 | 0.0107 | 0.0136 | 0.0091 | 0.0173 | 0.0118 |
| 1650 | uniform | 0.0203 | 0.0092 | 0.0415 | 0.0546 | 0.0345 | 0.0392 | 0.0252 |
| 3300 | uniform | 0.0155 | 0.0065 | 0.0301 | 0.0408 | 0.0260 | 0.0301 | 0.0180 |
| 6600 | uniform | 0.0109 | 0.0048 | 0.0221 | 0.0295 | 0.0177 | 0.0207 | 0.0135 |
| 9900 | uniform | 0.0081 | 0.0039 | 0.0180 | 0.0241 | 0.0133 | 0.0178 | 0.0102 |
| 13200 | uniform | 0.0070 | 0.0035 | 0.0159 | 0.0216 | 0.0125 | 0.0167 | 0.0086 |
| 19800 | uniform | 0.0056 | 0.0027 | 0.0123 | 0.0164 | 0.0096 | 0.0108 | 0.0077 |
| 1650 | population | 0.0197 | 0.0094 | 0.0427 | 0.0569 | 0.0358 | 0.0406 | 0.0235 |
| 3300 | population | 0.0151 | 0.0069 | 0.0316 | 0.0422 | 0.0260 | 0.0275 | 0.0167 |
| 6600 | population | 0.0101 | 0.0049 | 0.0222 | 0.0302 | 0.0178 | 0.0203 | 0.0126 |
| 9900 | population | 0.0086 | 0.0040 | 0.0182 | 0.0248 | 0.0147 | 0.0162 | 0.0103 |
| 13200 | population | 0.0070 | 0.0038 | 0.0175 | 0.0237 | 0.0128 | 0.0158 | 0.0086 |
| 19800 | population | 0.0057 | 0.0030 | 0.0137 | 0.0185 | 0.0102 | 0.0106 | 0.0078 |

| Size | Structure | OverallFoi | Foi1 | Foi2 | Foi3 | Foi4 | Foi5 | Foi6 |
|---|---|---|---|---|---|---|---|---|
| 1650 | survey | 0.0081 | 0.0169 | 0.0188 | 0.0160 | 0.0017 | 0.0048 | 0.0076 |
| 3300 | survey | 0.0064 | 0.0131 | 0.0146 | 0.0127 | 0.0014 | 0.0036 | 0.0060 |
| 6600 | survey | 0.0042 | 0.0079 | 0.0088 | 0.0076 | 0.0008 | 0.0022 | 0.0042 |
| 9900 | survey | 0.0039 | 0.0070 | 0.0078 | 0.0068 | 0.0007 | 0.0019 | 0.0040 |
| 13200 | survey | 0.0035 | 0.0064 | 0.0072 | 0.0062 | 0.0006 | 0.0018 | 0.0036 |
| 19800 | survey | 0.0025 | 0.0052 | 0.0058 | 0.0050 | 0.0005 | 0.0015 | 0.0025 |
| 1650 | uniform | 0.0095 | 0.0198 | 0.0220 | 0.0188 | 0.0020 | 0.0055 | 0.0079 |
| 3300 | uniform | 0.0068 | 0.0140 | 0.0155 | 0.0134 | 0.0014 | 0.0038 | 0.0059 |
| 6600 | uniform | 0.0051 | 0.0103 | 0.0114 | 0.0099 | 0.0011 | 0.0028 | 0.0045 |
| 9900 | uniform | 0.0042 | 0.0085 | 0.0095 | 0.0083 | 0.0009 | 0.0024 | 0.0039 |
| 13200 | uniform | 0.0038 | 0.0077 | 0.0086 | 0.0074 | 0.0008 | 0.0022 | 0.0037 |
| 19800 | uniform | 0.0028 | 0.0058 | 0.0065 | 0.0056 | 0.0006 | 0.0016 | 0.0024 |
| 1650 | population | 0.0099 | 0.0203 | 0.0225 | 0.0193 | 0.0020 | 0.0056 | 0.0085 |
| 3300 | population | 0.0072 | 0.0149 | 0.0165 | 0.0142 | 0.0015 | 0.0041 | 0.0061 |
| 6600 | population | 0.0053 | 0.0107 | 0.0118 | 0.0102 | 0.0011 | 0.0030 | 0.0047 |
| 9900 | population | 0.0042 | 0.0086 | 0.0096 | 0.0083 | 0.0009 | 0.0024 | 0.0035 |
| 13200 | population | 0.0041 | 0.0083 | 0.0093 | 0.0081 | 0.0009 | 0.0023 | 0.0037 |
| 19800 | population | 0.0031 | 0.0064 | 0.0072 | 0.0062 | 0.0007 | 0.0018 | 0.0026 |

| **Size** | **Structure** | **$R_0$** | **$R_{eff}$** | **ASI** | **$\varphi$** |
|---|---|---|---|---|---|
| 1650 | survey | 0.4543 | 0.0088 | 1.2815 | 1.0859 |
| 3300 | survey | 0.3533 | 0.0069 | 1.0170 | 0.4921 |
| 6600 | survey | 0.2126 | 0.0042 | 0.6099 | 0.3999 |
| 9900 | survey | 0.1882 | 0.0037 | 0.5419 | 0.3306 |
| 13200 | survey | 0.1732 | 0.0034 | 0.4963 | 0.3162 |
| 19800 | survey | 0.1406 | 0.0027 | 0.3992 | 0.1803 |
| 1650 | uniform | 0.5320 | 0.0103 | 1.5003 | 0.4664 |
| 3300 | uniform | 0.3756 | 0.0073 | 1.0588 | 0.3797 |
| 6600 | uniform | 0.2757 | 0.0054 | 0.7776 | 0.2191 |
| 9900 | uniform | 0.2298 | 0.0045 | 0.6626 | 0.2534 |
| 13200 | uniform | 0.2067 | 0.0040 | 0.5953 | 0.1764 |
| 19800 | uniform | 0.1573 | 0.0031 | 0.4478 | 0.0994 |
| 1650 | population | 0.5448 | 0.0106 | 1.5179 | 0.5419 |
| 3300 | population | 0.3988 | 0.0078 | 1.1245 | 0.3904 |
| 6600 | population | 0.2861 | 0.0056 | 0.8054 | 0.1948 |
| 9900 | population | 0.2315 | 0.0045 | 0.6633 | 0.1386 |
| 13200 | population | 0.2245 | 0.0044 | 0.6458 | 0.1613 |
| 19800 | population | 0.1735 | 0.0034 | 0.4946 | 0.0994 |

### Optimal distributions

In Tables S13-20, the optimal distributions of samples for the overall and age-specific seroprevalence and force of infection by pathogen, model, and sample size are given. The age groups (Group 1-6) are ([1,2), [2,6), [6,12), [12,19), [19,31), and [31,65] years.

**Table S13.** Measles serological data – Logistic model with piecewise constant prevalence

| Size | Group 1 % | Group 2 % | Group 3 % | Group 4 % | Group 5 % | Group 6 % | Precision |
|---|---|---|---|---|---|---|---|
| **Overall prevalence** | | | | | | | |
| 1650 | 10 | 10 | 10 | 20 | 20 | 30 | 0.0104 |
| 3300 | 10 | 20 | 10 | 10 | 20 | 30 | 0.0081 |
| 6600 | 10 | 10 | 10 | 20 | 20 | 30 | 0.0053 |
| 9900 | 10 | 10 | 10 | 10 | 30 | 30 | 0.0042 |
| 13200 | 10 | 10 | 10 | 20 | 20 | 30 | 0.0036 |
| 19800 | 10 | 10 | 10 | 20 | 20 | 30 | 0.0031 |
| **Prevalence by age group** | | | | | | | |
| 1650 | 20 | 20 | 10 | 20 | 20 | 10 | 0.1950 |
| 3300 | 30 | 10 | 20 | 10 | 20 | 10 | 0.1332 |
| 6600 | 20 | 20 | 10 | 20 | 20 | 10 | 0.0953 |
| 9900 | 20 | 20 | 10 | 20 | 20 | 10 | 0.0759 |
| 13200 | 30 | 10 | 20 | 10 | 20 | 10 | 0.0675 |
| 19800 | 20 | 10 | 20 | 10 | 20 | 20 | 0.0567 |

**Table S14.** Mumps serological data – Logistic model with piecewise constant prevalence

| Size | Group 1 % | Group 2 % | Group 3 % | Group 4 % | Group 5 % | Group 6 % | Precision |
|---|---|---|---|---|---|---|---|
| **Overall prevalence** | | | | | | | |
| 1650 | 10 | 10 | 10 | 10 | 10 | 50 | 0.0191 |
| 3300 | 10 | 20 | 10 | 10 | 10 | 40 | 0.0143 |
| 6600 | 10 | 10 | 10 | 10 | 10 | 50 | 0.0097 |
| 9900 | 10 | 10 | 10 | 10 | 20 | 40 | 0.0080 |
| 13200 | 10 | 10 | 10 | 10 | 10 | 50 | 0.0070 |
| 19800 | 10 | 10 | 10 | 20 | 10 | 40 | 0.0062 |
| **Prevalence by age group** | | | | | | | |
| 1650 | 20 | 20 | 10 | 10 | 20 | 20 | 0.2642 |
| 3300 | 20 | 10 | 30 | 10 | 20 | 10 | 0.1877 |
| 6600 | 20 | 20 | 10 | 10 | 20 | 20 | 0.1289 |
| 9900 | 20 | 10 | 20 | 10 | 20 | 20 | 0.1041 |
| 13200 | 20 | 10 | 20 | 10 | 20 | 20 | 0.0916 |
| 19800 | 20 | 20 | 10 | 10 | 20 | 20 | 0.0752 |

**Table S15.** Rubella serological data – Logistic model with piecewise constant prevalence

| Size | Group 1 % | Group 2 % | Group 3 % | Group 4 % | Group 5 % | Group 6 % | Precision |
|---|---|---|---|---|---|---|---|
| **Overall prevalence** | | | | | | | |
| 1650 | 10 | 10 | 10 | 20 | 10 | 40 | 0.0106 |
| 3300 | 10 | 10 | 10 | 20 | 10 | 40 | 0.0077 |
| 6600 | 10 | 10 | 10 | 10 | 20 | 40 | 0.0054 |
| 9900 | 10 | 10 | 10 | 10 | 30 | 30 | 0.0044 |
| 13200 | 10 | 10 | 10 | 10 | 20 | 40 | 0.0037 |
| 19800 | 10 | 10 | 10 | 20 | 10 | 40 | 0.0031 |
| **Prevalence by age group** | | | | | | | |
| 1650 | 20 | 20 | 20 | 10 | 10 | 20 | 0.1793 |
| 3300 | 30 | 10 | 20 | 10 | 20 | 10 | 0.1269 |
| 6600 | 20 | 20 | 20 | 10 | 10 | 20 | 0.0873 |
| 9900 | 30 | 20 | 10 | 10 | 10 | 20 | 0.0735 |
| 13200 | 30 | 10 | 30 | 10 | 10 | 10 | 0.0635 |
| 19800 | 30 | 10 | 20 | 10 | 20 | 10 | 0.0510 |

**Table S16.** VZV serological data – Exponentially damped model

| Size | Group 1 % | Group 2 % | Group 3 % | Group 4 % | Group 5 % | Group 6 % | Precision |
|---|---|---|---|---|---|---|---|
| **Overall prevalence** | | | | | | | |
| 1650 | 30 | 10 | 10 | 30 | 10 | 10 | 0.0048 |
| 3300 | 10 | 30 | 10 | 10 | 30 | 10 | 0.0050 |
| 6600 | 10 | 10 | 10 | 10 | 40 | 20 | 0.0020 |
| 9900 | 10 | 10 | 10 | 10 | 40 | 20 | 0.0021 |
| 13200 | 50 | 10 | 10 | 10 | 10 | 10 | 0.0020 |
| 19800 | 10 | 10 | 20 | 10 | 30 | 20 | 0.0019 |
| **Prevalence by age group** | | | | | | | |
| 1650 | 20 | 20 | 20 | 20 | 10 | 10 | 0.1081 |
| 3300 | 20 | 40 | 10 | 10 | 10 | 10 | 0.0859 |
| 6600 | 10 | 20 | 10 | 40 | 10 | 10 | 0.0585 |
| 9900 | 10 | 20 | 10 | 10 | 30 | 20 | 0.0520 |
| 13200 | 10 | 30 | 20 | 10 | 10 | 20 | 0.0420 |
| 19800 | 20 | 20 | 30 | 10 | 10 | 10 | 0.0342 |
| **Overall force of infection** | | | | | | | |
| 1650 | 20 | 20 | 20 | 10 | 20 | 10 | 0.0264 |
| 3300 | 10 | 30 | 10 | 10 | 30 | 10 | 0.0255 |
| 6600 | 10 | 10 | 10 | 10 | 40 | 20 | 0.0094 |
| 9900 | 10 | 10 | 10 | 10 | 40 | 20 | 0.0101 |
| 13200 | 50 | 10 | 10 | 10 | 10 | 10 | 0.0096 |
| 19800 | 10 | 10 | 10 | 20 | 30 | 20 | 0.0096 |
| **Force of infection by age group** | | | | | | | |
| 1650 | 10 | 20 | 10 | 10 | 30 | 20 | 0.2593 |
| 3300 | 10 | 20 | 20 | 10 | 20 | 20 | 0.1842 |
| 6600 | 10 | 20 | 20 | 20 | 10 | 20 | 0.1208 |
| 9900 | 10 | 20 | 10 | 10 | 10 | 40 | 0.0991 |
| 13200 | 10 | 30 | 20 | 10 | 10 | 20 | 0.0883 |
| 19800 | 20 | 20 | 20 | 10 | 10 | 20 | 0.0696 |

**Table S17.** VZV serological data – MSIR piecewise constant force of infection

| Size | Group 1 % | Group 2 % | Group 3 % | Group 4 % | Group 5 % | Group 6 % | Precision |
|---|---|---|---|---|---|---|---|
| **Overall prevalence** | | | | | | | |
| 1650 | 10 | 20 | 10 | 10 | 30 | 20 | 0.0069 |
| 3300 | 10 | 20 | 20 | 10 | 30 | 10 | 0.0048 |
| 6600 | 10 | 20 | 10 | 10 | 30 | 20 | 0.0032 |
| 9900 | 10 | 10 | 20 | 10 | 30 | 20 | 0.0026 |
| 13200 | 10 | 10 | 20 | 10 | 30 | 20 | 0.0024 |
| 19800 | 10 | 10 | 20 | 10 | 30 | 20 | 0.0020 |
| **Prevalence by age group** | | | | | | | |
| 1650 | 10 | 40 | 10 | 20 | 10 | 10 | 0.1229 |
| 3300 | 20 | 30 | 20 | 10 | 10 | 10 | 0.0914 |
| 6600 | 10 | 40 | 10 | 20 | 10 | 10 | 0.0630 |
| 9900 | 10 | 50 | 10 | 10 | 10 | 10 | 0.0538 |
| 13200 | 10 | 30 | 20 | 10 | 20 | 10 | 0.0464 |
| 19800 | 10 | 40 | 20 | 10 | 10 | 10 | 0.0374 |
| **Overall force of infection** | | | | | | | |
| 1650 | 10 | 10 | 40 | 10 | 10 | 20 | 0.0356 |
| 3300 | 10 | 10 | 30 | 10 | 30 | 10 | 0.0266 |
| 6600 | 10 | 10 | 30 | 10 | 20 | 20 | 0.0187 |
| 9900 | 10 | 10 | 20 | 10 | 20 | 30 | 0.0144 |
| 13200 | 10 | 10 | 20 | 10 | 30 | 20 | 0.0132 |
| 19800 | 10 | 10 | 20 | 10 | 30 | 20 | 0.0106 |
| **Force of infection by age group** | | | | | | | |
| 1650 | 10 | 40 | 10 | 10 | 20 | 10 | 0.5954 |
| 3300 | 10 | 30 | 10 | 30 | 10 | 10 | 0.5147 |
| 6600 | 10 | 20 | 10 | 10 | 10 | 40 | 0.3494 |
| 9900 | 10 | 10 | 20 | 10 | 10 | 40 | 0.2953 |
| 13200 | 10 | 10 | 20 | 20 | 10 | 30 | 0.2529 |
| 19800 | 10 | 10 | 20 | 10 | 20 | 30 | 0.1976 |

**Table S18.** Parvovirus B19 serological data – Exponentially damped model

| Size | Group 1 % | Group 2 % | Group 3 % | Group 4 % | Group 5 % | Group 6 % | Precision |
|---|---|---|---|---|---|---|---|
| **Overall prevalence** | | | | | | | |
| 1650 | 10 | 10 | 20 | 10 | 20 | 30 | 0.0214 |
| 3300 | 10 | 10 | 10 | 20 | 10 | 40 | 0.0146 |
| 6600 | 10 | 10 | 10 | 20 | 10 | 40 | 0.0107 |
| 9900 | 10 | 10 | 10 | 10 | 20 | 40 | 0.0088 |
| 13200 | 20 | 10 | 10 | 10 | 20 | 30 | 0.0074 |
| 19800 | 10 | 10 | 10 | 10 | 20 | 40 | 0.0063 |
| **Prevalence by age group** | | | | | | | |
| 1650 | 10 | 20 | 10 | 20 | 20 | 20 | 0.1725 |
| 3300 | 10 | 20 | 10 | 20 | 10 | 30 | 0.1258 |
| 6600 | 10 | 10 | 30 | 10 | 20 | 20 | 0.0888 |
| 9900 | 10 | 30 | 10 | 20 | 10 | 20 | 0.0748 |
| 13200 | 10 | 20 | 20 | 10 | 10 | 30 | 0.0634 |
| 19800 | 10 | 20 | 20 | 10 | 20 | 20 | 0.0536 |
| **Overall force of infection** | | | | | | | |
| 1650 | 10 | 10 | 10 | 10 | 10 | 50 | 0.0043 |
| 3300 | 10 | 10 | 10 | 10 | 10 | 50 | 0.0030 |
| 6600 | 10 | 10 | 10 | 10 | 10 | 50 | 0.0021 |
| 9900 | 10 | 10 | 10 | 10 | 10 | 50 | 0.0018 |
| 13200 | 10 | 10 | 10 | 10 | 10 | 50 | 0.0015 |
| 19800 | 10 | 10 | 10 | 10 | 10 | 50 | 0.0014 |
| **Force of infection by age group** | | | | | | | |
| 1650 | 10 | 30 | 10 | 20 | 10 | 20 | 0.0630 |
| 3300 | 10 | 30 | 10 | 20 | 10 | 20 | 0.0458 |
| 6600 | 10 | 40 | 10 | 10 | 10 | 20 | 0.0329 |
| 9900 | 10 | 30 | 10 | 10 | 20 | 20 | 0.0264 |
| 13200 | 10 | 30 | 10 | 10 | 20 | 20 | 0.0232 |
| 19800 | 10 | 20 | 20 | 10 | 20 | 20 | 0.0195 |

**Table S19.** Parvovirus B19 serological data – MSIR piecewise constant force of infection

| Size | Group 1 % | Group 2 % | Group 3 % | Group 4 % | Group 5 % | Group 6 % | Precision |
|---|---|---|---|---|---|---|---|
| **Overall prevalence** | | | | | | | |
| 1650 | 10 | 10 | 20 | 10 | 20 | 30 | 0.0212 |
| 3300 | 10 | 10 | 10 | 10 | 20 | 40 | 0.0147 |
| 6600 | 10 | 10 | 10 | 10 | 30 | 30 | 0.0106 |
| 9900 | 10 | 10 | 10 | 10 | 20 | 40 | 0.0087 |
| 13200 | 10 | 10 | 10 | 10 | 20 | 40 | 0.0072 |
| 19800 | 10 | 10 | 10 | 10 | 10 | 50 | 0.0064 |
| **Prevalence by age group** | | | | | | | |
| 1650 | 10 | 20 | 20 | 10 | 20 | 20 | 0.2059 |
| 3300 | 10 | 10 | 20 | 10 | 20 | 30 | 0.1527 |
| 6600 | 10 | 20 | 20 | 20 | 10 | 20 | 0.1086 |
| 9900 | 10 | 30 | 10 | 20 | 10 | 20 | 0.0869 |
| 13200 | 10 | 20 | 20 | 10 | 20 | 20 | 0.0754 |
| 19800 | 10 | 20 | 20 | 20 | 10 | 20 | 0.0628 |
| **Overall force of infection** | | | | | | | |
| 1650 | 10 | 10 | 10 | 10 | 10 | 50 | 0.0047 |
| 3300 | 10 | 10 | 10 | 10 | 10 | 50 | 0.0033 |
| 6600 | 10 | 10 | 10 | 10 | 10 | 50 | 0.0022 |
| 9900 | 10 | 10 | 10 | 10 | 10 | 50 | 0.0020 |
| 13200 | 10 | 10 | 10 | 10 | 10 | 50 | 0.0017 |
| 19800 | 10 | 10 | 10 | 10 | 10 | 50 | 0.0015 |
| **Force of infection by age group** | | | | | | | |
| 1650 | 10 | 40 | 10 | 10 | 10 | 20 | 0.1692 |
| 3300 | 20 | 20 | 10 | 20 | 10 | 20 | 0.1299 |
| 6600 | 10 | 20 | 20 | 10 | 20 | 20 | 0.0936 |
| 9900 | 10 | 20 | 30 | 10 | 20 | 10 | 0.0762 |
| 13200 | 10 | 20 | 40 | 10 | 10 | 10 | 0.0669 |
| 19800 | 10 | 20 | 30 | 10 | 10 | 20 | 0.0536 |

**Table S20.** Parvovirus B19 serological data – MSIRWb-ext AW model

| Size | Group 1 % | Group 2 % | Group 3 % | Group 4 % | Group 5 % | Group 6 % | Precision |
|---|---|---|---|---|---|---|---|
| **Overall prevalence** | | | | | | | |
| 1650 | 10 | 10 | 10 | 20 | 20 | 30 | 0.0237 |
| 3300 | 10 | 10 | 10 | 10 | 10 | 50 | 0.0156 |
| 6600 | 10 | 10 | 10 | 10 | 20 | 40 | 0.0115 |
| 9900 | 10 | 10 | 10 | 10 | 20 | 40 | 0.0090 |
| 13200 | 10 | 10 | 10 | 10 | 20 | 40 | 0.0075 |
| 19800 | 10 | 10 | 10 | 10 | 10 | 50 | 0.0067 |
| **Prevalence by age group** | | | | | | | |
| 1650 | 10 | 10 | 20 | 20 | 20 | 20 | 0.1921 |
| 3300 | 10 | 10 | 20 | 10 | 10 | 40 | 0.1435 |
| 6600 | 10 | 10 | 10 | 20 | 20 | 30 | 0.1029 |
| 9900 | 10 | 10 | 20 | 10 | 10 | 40 | 0.0817 |
| 13200 | 10 | 10 | 20 | 10 | 10 | 40 | 0.0740 |
| 19800 | 10 | 20 | 20 | 10 | 10 | 30 | 0.0615 |
| **Overall force of infection** | | | | | | | |
| 1650 | 10 | 20 | 20 | 20 | 10 | 20 | 0.0076 |
| 3300 | 10 | 10 | 30 | 10 | 20 | 20 | 0.0058 |
| 6600 | 10 | 20 | 20 | 10 | 10 | 30 | 0.0042 |
| 9900 | 10 | 10 | 20 | 10 | 10 | 40 | 0.0035 |
| 13200 | 10 | 10 | 20 | 10 | 10 | 40 | 0.0030 |
| 19800 | 10 | 10 | 20 | 30 | 10 | 20 | 0.0023 |
| **Force of infection by age group** | | | | | | | |
| 1650 | 10 | 10 | 30 | 20 | 10 | 20 | 0.0596 |
| 3300 | 10 | 10 | 30 | 10 | 20 | 20 | 0.0444 |
| 6600 | 10 | 20 | 20 | 10 | 10 | 30 | 0.0339 |
| 9900 | 10 | 10 | 20 | 20 | 10 | 30 | 0.0265 |
| 13200 | 10 | 10 | 20 | 10 | 10 | 40 | 0.0245 |
| 19800 | 10 | 10 | 20 | 30 | 10 | 20 | 0.0184 |
